# Proteasomal subunit depletions differentially affect germline integrity in *C. elegans*

**DOI:** 10.1101/2022.03.21.485201

**Authors:** Lourds Michelle Fernando, Cristina Quesada-Candela, Makaelah Murray, Caroline Ugoaru, Judith L. Yanowitz, Anna K. Allen

**Author notes:** **Correspondence:** Anna K. Allen, Judith L. Yanowitz. These authors have contributed equally to this work and share first authorship (Lourds Michelle Fernando, Cristina Quesada-Candela).

## Abstract

The 26S proteasome is a multi-subunit protein complex that is canonically known for its ability to degrade proteins in cells and maintain protein homeostasis. Non-canonical or non-proteolytic roles of proteasomal subunits exist, but remain less well studied. We provide characterization of germline-specific functions of different 19S RP proteasome subunits in *C. elegans* using RNAi specifically from the L4 stage and through generation of endogenously tagged 19S RP lid subunit strains. We show functions for the 19S RP in regulation of proliferation and maintenance of integrity of mitotic zone nuclei, in polymerization of the synaptonemal complex (SC) onto meiotic chromosomes and in the timing of SC subunit redistribution to the short arm of the bivalent, and in turnover of XND-1 proteins at late pachytene. Furthermore, we report that certain 19S RP subunits are required for proper germ line localization of WEE-1.3, a major meiotic kinase. Additionally, endogenous fluorescent labeling revealed that the two isoforms of the essential 19S RP proteasome subunit RPN-6.1 are expressed in a tissue-specific manner in the hermaphrodite. Also, we demonstrate that the 19S RP subunits RPN-6.1 and RPN-7 are crucial for the nuclear localization of the lid subunits RPN-8 and RPN-9 in oocytes, potentially introducing *C. elegans* germ line as model to study proteasome assembly real-time. Collectively, our data support the premise that certain 19S RP proteasome subunits are playing tissue-specific roles, especially in the germ line. We propose *C. elegans* as a versatile multicellular model to study the diverse proteolytic and non-proteolytic roles that proteasome subunits play *in vivo*.

## Introduction

The 26S proteasome is a ~2.5 MDa multi-subunit protein complex that maintains cellular homeostasis by degrading old, misfolded, mistranslated, and/or regulatory proteins in cells in both the cytoplasm and the nucleus (Hanna and Finley, 2007; Pack *et al.*, 2014; Bard *et al.*, 2018; Marshall and Vierstra, 2019). Recent evidence shows that specific proteasome subunits play tissue specific and/or non-proteolytic roles in various organisms (Pispa *et al.*, 2008; Bhat and Greer, 2011; Pispa, Matilainen and Holmberg, 2020). This includes roles in various cellular processes such as transcription, mRNA export, cell cycle regulation and chromosome structure maintenance (Ferdous, Kodadek and Johnston, 2002; Kwak, Workman and Lee, 2011; Seo *et al.*, 2017; Gómez-H *et al.*, 2019). Models such as yeast and mammalian cell lines are widely used to characterize proteasome function, however, these unicellular models have limitations in comprehensively understanding the wide range of roles that individual proteasome subunits might be playing in different tissues and developmental stages (Hochstrasser, 1996; Bai *et al.*, 2019). Proper understanding of the assembly, structure, and function of the proteasome is crucial for understanding the pathology of diseases caused by irregular proteasome function, such as neurodegenerative diseases and cancer(Hanna and Finley, 2007; Hirano *et al.*, 2008; Myeku *et al.*, 2011; Kish-Trier and Hill, 2013; Saez and Vilchez, 2014; Schmidt and Finley, 2014; Maneix and Catic, 2016; Walerych *et al.*, 2016).

High resolution structural characterization of the 26S proteasome in human and yeast via cryo-electron microscopy and atomic modeling has revealed the structure of the eukaryotic proteasome at atomic level (Groll *et al.*, 1997; Unno *et al.*, 2002; Beck *et al.*, 2012; Li *et al.*, 2013; Huang *et al.*, 2016). The mature 26S proteasome is composed of approximately 33 different, highly conserved protein subunits arranged into two 19S regulatory particles (RP) capping one cylindrical 20S core particle (CP) (Figure 1A) (Kish-Trier and Hill, 2013). The 20S CP possesses the peptidase activity to degrade a protein substrate into smaller peptides, while the 19S RPs are responsible for recognizing, deubiquitinating and unfolding of polyubiquitinated substrates before importing substrates into the CP (Hanna and Finley, 2007; Finley, 2009). Each 19S RP is made up of two sub-complexes referred to as the lid and the base. The 19S RP lid is composed of non-ATPase subunits (Rpn3, Rpn5, Rpn6, Rpn7, Rpn8, Rpn9, Rpn11, Rpn12 and Sem1), while the base is composed of three non-ATPase subunits (Rpn1, Rpn2, and Rpn13) and six ATPase subunits (Rpt1, Rpt2, Rpt3, Rpt4, Rpt5, and Rpt6) (Kim, Yu and Cheng, 2011; Uprety *et al.*, 2012). A final subunit, Rpn10, is thought to bridge the lid and base subcomplexes thus joining the two together (Bard *et al.*, 2018). The *C. elegans* proteins comprising the 26S proteasome are diagrammed in Figure 1A and listed along with their human and yeast orthologs in Supplemental Table 1.

**Figure 1.**
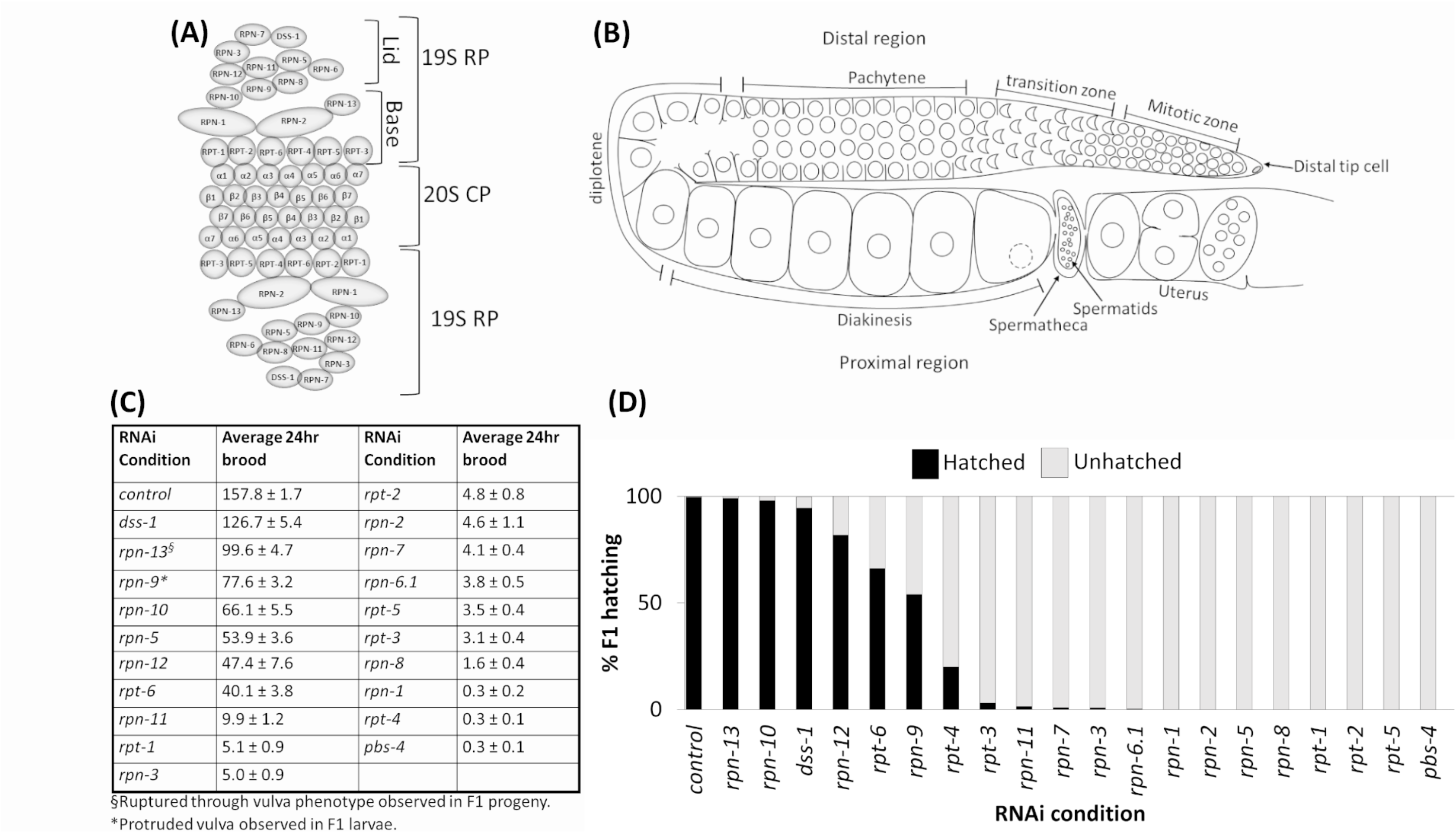
Depletion of 19S RP subunits of the 26S proteasome in *C. elegans* hermaphrodites caused reduced brood and/or embryonic lethality. (A) Schematic of eukaryotic 26S proteasome and its subunits. (B) Schematic of an adult *C. elegans* hermaphrodite germ line (one gonad arm). (C) Average 24 hr brood of *C. elegans* hermaphrodites RNAi-depleted of either a control gene (n=152), any of the 19 subunits of the 19S RP (n=10-83), or a 20S CP subunit, PBS-4 (n=36). Brood is shown ± SEM and calculated from at least three independent trials. All RNAi conditions compared to control exhibit a p-value < 0.0001. (D) Percent of hatched (black bars) and unhatched (grey bars) progeny of hermaphrodites treated with either *control(RNAi)* or the indicated *proteasome subunit(RNAi)*.

Assembly of the subunits to make a functional 26S proteasome is a highly conserved, multistep process. The 20S CP and 19S RP assemble independently as subcomplexes in the cytoplasm and then either can combine into the 26S in this compartment or can be imported into the nucleus and then assemble to form the mature 26S structure (Hirano *et al.*, 2006; Kusmierczyk *et al.*, 2008; Pack *et al.*, 2014; Budenholzer *et al.*, 2017; Marshall and Vierstra, 2019). The 20S CP subcomplex assembly is known to require the aid of non-proteasomal chaperone proteins, while nuclear localization sequences (NLSs) on the alpha subunits of the 20S CP aid in the nuclear import of the subcomplexes (Brooks *et al.*, 2000; Hirano *et al.*, 2006; Kusmierczyk *et al.*, 2008; Budenholzer *et al.*, 2017; Wu *et al.*, 2018). The 19S RP lid and base subcomplexes assemble separately in the cytoplasm, before being imported into the nucleus where the separate modules dock on the assembled 20S CP to form the mature 26S proteasome (Tanaka *et al.*, 1990; Lehmann *et al.*, 2002; Wendler *et al.*, 2004). Previous research in yeast has identified assembly chaperones for the 19S RP base subcomplex and NLSs on two base subunits (yeast Rpt2 and Rpn2) aid in the nuclear import of the base (Wendler *et al.*, 2004; Wendler and Enenkel, 2019). The yeast 19S RP lid subcomplex assembly consists first of the formation of Module 1 (Rpn5, Rpn6, Rpn8, Rpn9 and Rpn11) which then binds to lid particle 3 (Rpn3, Rpn7 and Sem1/Dss1) with Rpn12 serving as the linker (Budenholzer *et al.*, 2017). Interestingly, no external factors or assembly chaperones have yet been identified that assist in 19S RP lid subcomplex assembly, nor do any of the lid subcomplex proteins have known NLS sequences which could aid in the nuclear import of the 19S lid (Isono *et al.*, 2007; Budenholzer *et al.*, 2020). Therefore, further studies are required to fill the gap in our understanding of nuclear import of the 19S lid subcomplex.

While the role of the proteasome as the protein degradation machine in eukaryotes is well characterized, recent findings have sparked an interest in non-canonical and tissue-specific roles of individual proteasome subunits and/or subcomplexes. In mammals, tissue-specific proteasomes, such as the immunoproteasome, thymoproteasome, and spermatoproteasome contain structural variations in specific proteasome subunits leading to their tissue specificity (Kish-Trier and Hill, 2013; Uechi, Hamazaki and Murata, 2014; Gómez-H *et al.*, 2019; Motosugi and Murata, 2019). Studies done in mammals and *C. elegans* show that the 19S RP lid subunit PSMD11/RPN-6.1 can regulate proteolytic activity of the proteasome modulating the production of the other proteasome subunits thus increasing or decreasing proteolytic activity of the proteasome (Vilchez, Boyer, *et al.*, 2012; Vilchez, Morantte, *et al.*, 2012; Lokireddy, Kukushkin and Goldberg, 2015). *C. elegans* studies have also uncovered proteasome subunits that are specific for germline development and fertility (Shimada *et al.*, 2006; Pispa *et al.*, 2008; Fernando, Elliot and Allen, 2020). RPN-10, RPN-12 and DSS-1 (RPN15/SEM1) were each shown to play specific roles in germline sex determination and oocyte development (Shimada *et al.*, 2006; Pispa *et al.*, 2008; Fernando, Elliot and Allen, 2020).

Proper function of the 26S proteasome in the *C. elegans* hermaphrodite germ line is crucial for normal progression of meiosis and production of viable progeny (Glotzer, Murray and Kirschner, 1991; Lee and Schedl, 2010). The two germ line arms of the nematode meet at a shared uterus. Each arm contains a distal mitotic pool of cells that enter meiosis as they move proximally (Figure 1B) (Hubbard and Greenstein, 2000; Hillers *et al.*, 2015). The germ line nuclei are open to the central rachis until the diplotene stage when cellularization of the developing oocytes is completed. The oocytes briefly arrest at the diakinesis stage prior to maturation, ovulation, and completion of the meiotic divisions (Greenstein, 2005). Feeding L4 *C. elegans* hermaphrodites dsRNA against individual 19S RP proteasome subunits results in F1 progeny lethality for most of the 19S RP subunits, the exceptions being RPN-9, RPN-10, RPN-12, DSS-1, and RPT-6 (Takahashi *et al.*, 2002; Shimada *et al.*, 2006; Pispa *et al.*, 2008; Fernando, Elliot and Allen, 2020). Despite the impact on embryonic viability, the effect of 19S RP subunit depletion on the reproductive capabilities of the RNAi-treated hermaphrodite mothers has not been examined. Here we report fertility defects observed in *C. elegans* hermaphrodites RNAi-depleted of individual 19S RP subunits starting from the L4 stage. Our study includes testing of 19S RP subunits that were not part of a 2002 study that reported the embryonic lethality effect of RNAi depletion of various of the 26S proteasomal subunits (Takahashi *et al.*, 2002).

Recently our labs separately characterized previously unknown roles for the proteasome in the germ line (Allen, Nesmith and Golden, 2014; Ahuja *et al.*, 2017; Fernando, Elliot and Allen, 2020). We reported interactions between specific 19S RP subunits with a major meiotic kinase, WEE-1.3; we also described synaptonemal complex (SC) defects upon impairment of the 20S proteasome (Allen, Nesmith and Golden, 2014; Ahuja *et al.*, 2017; Fernando, Elliot and Allen, 2020). Here, we have embarked on a more detailed analysis of individual proteasomal subunit function in both the distal and proximal germ line of the *C. elegans* hermaphrodite. *C. elegans* is a powerful genetic model whose optical transparency enables the observation of biological processes in real-time and the determination of the subcellular localization of fluorescently tagged proteins of interest during any stage of the *C. elegans* life cycle. To help elucidate individual proteasome subunit functions in the germ line, we began endogenously tagging 19S RP lid subunits with GFP or OLLAS, and present novel tissue-specific expression of RPN-6.1 and genetic requirements for the nuclear localization of lid subunits RPN-8 and RPN-9 in the *C. elegans* oocyte. We propose *C. elegans* as a versatile multicellular model to study the diverse proteolytic and non-proteolytic roles proteasome subunits play *in vivo* in specific tissues and cell types.

## Materials Methods

### Strains

All strains were maintained at 20°C on standard MYOB or NGM plates seeded with OP50 unless mentioned otherwise (Brenner, 1974). Bristol strain N2 was used as the wild-type strain. Other strains used in this study are included in Supplemental Table 2.

### Strain generation

Strains in this study were generated using CRISPR/Cas9 genome editing technology following the direct delivery method developed by Paix *et al.* 2015 (Paix *et al.*, 2015). The Co-CRISPR method using *unc-58* or *dpy-10* was performed to screen for desired edits (Arribere *et al.*, 2014). Specificity of the crRNAs were determined using UCSC genome browser and http://crispr.mit.edu/. ApE plasmid editor was used for sequence analysis to select PAM sites and primer designs. The edits were confirmed using PCR. At least two independent strains were generated for each edit (except N-terminal GFP tagged RPN-7 for which only 1 strain was generated) and the resulting edited strains backcrossed with wild type (N2) at least 5 times and sequenced before being utilized.

GFP tags were generated by inserting Superfolder GFP sequence at the N-terminus immediately after the start ATG. Repair templates for the GFP strains were generated by PCR amplifying Superfolder GFP from pDONR221. All the strains generated in this study can be found in Table 2. The list of crRNAs (Horizon Discovery Ltd.) and primers (IDT Inc. or Eurofins genomics) used for generating repair templates and for PCR screening to confirm successful edits are listed in Supplemental Tables 3 and 4 respectively.

**Table 1:**
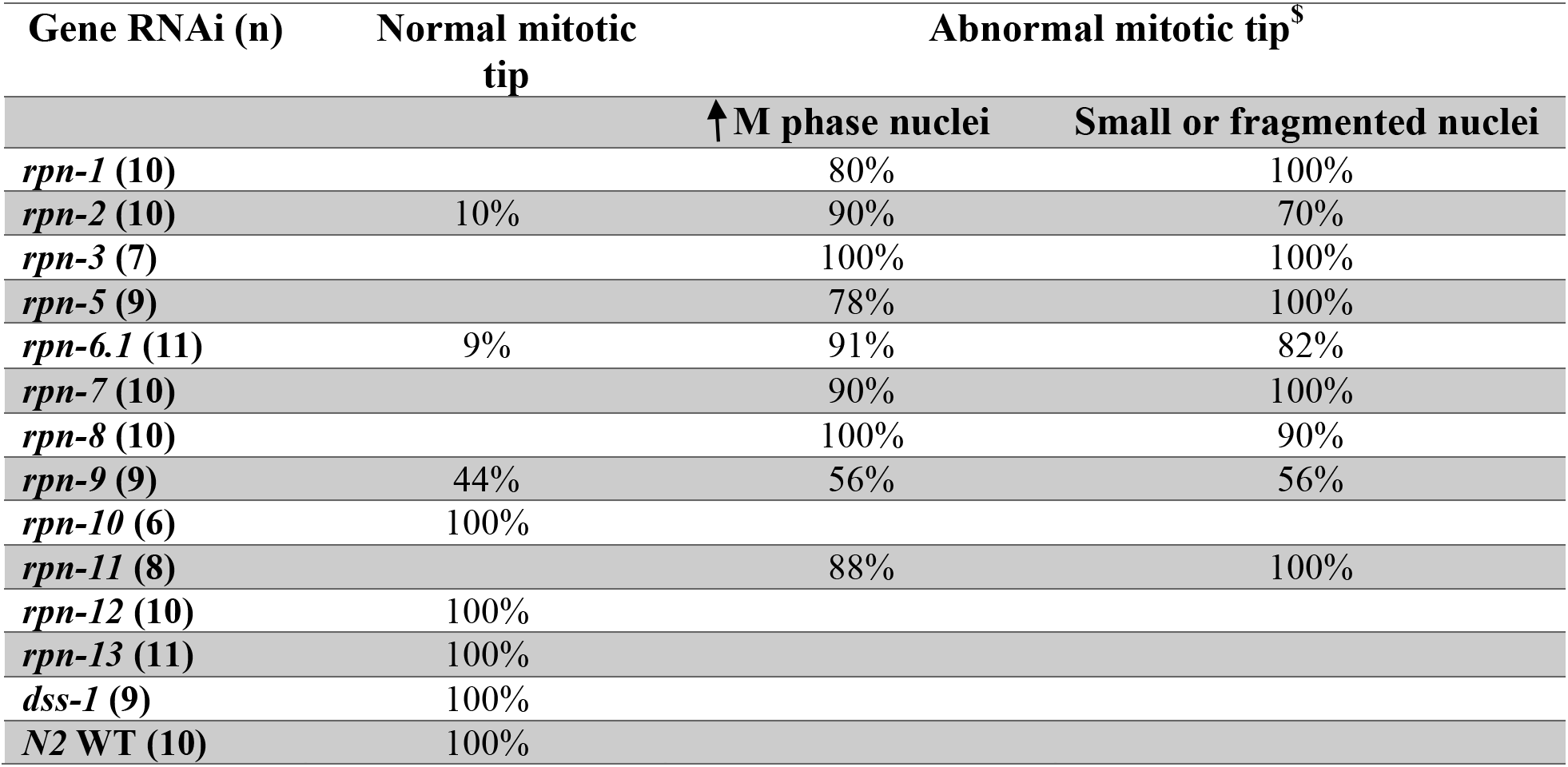
Percentage of worms that presented cell cycle defects after knocking down proteasome non-ATPase subunits.

**Table 2:**
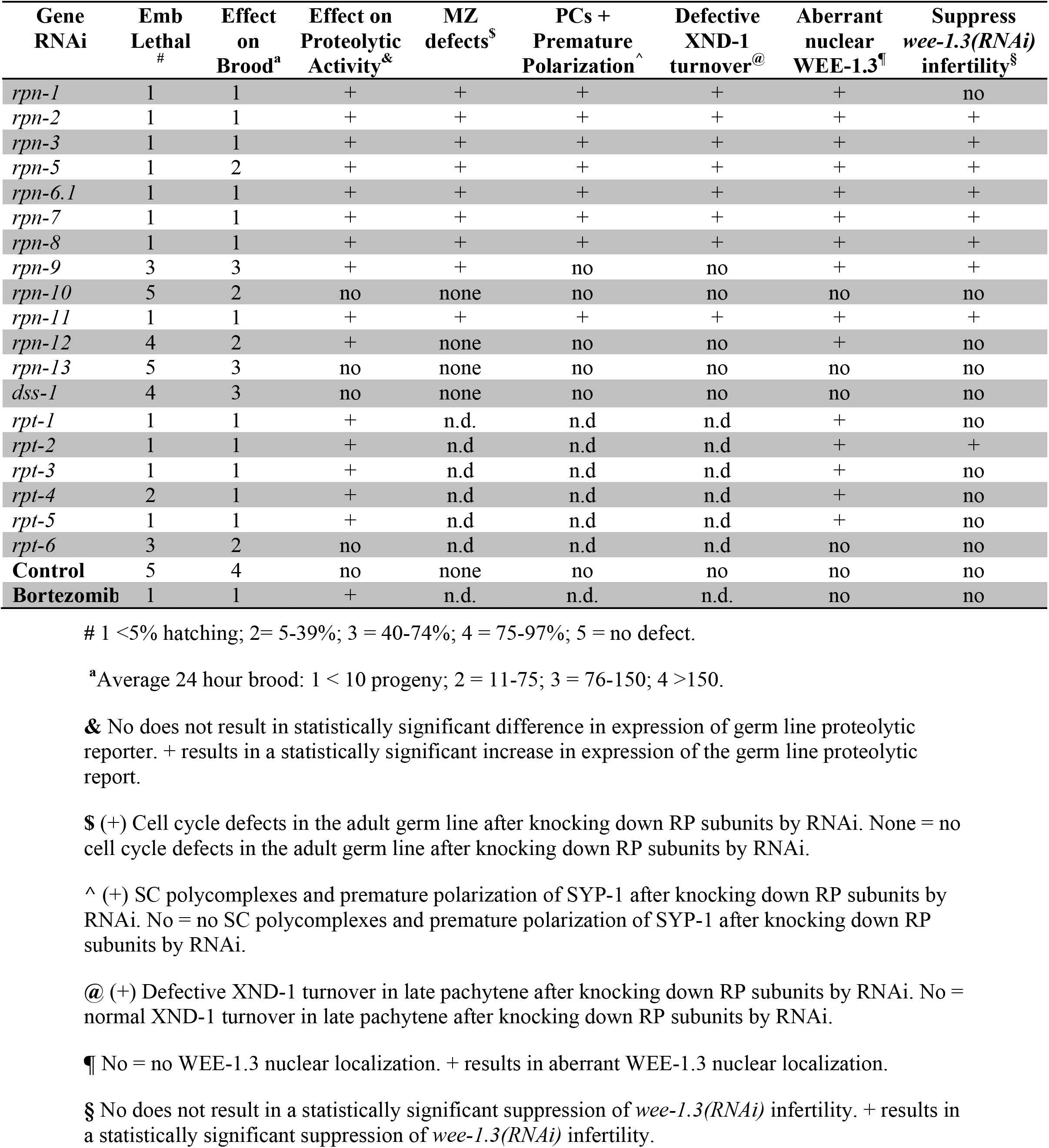
Summary of the germline phenotypes associated with RNAi-depletion of the various 19S RP subunits.

The C-terminal OLLAS-tag for RPN-6.1 was generated by inserting the 42bp OLLAS sequence, 5’-tccggattcgccaacgagctcggaccacgtctcatgggaaag-3’ immediately before the stop codon (TGA) in *rpn-6.1*. An ssODN was used as the repair template and contained a minimum of 35bp homology arms to the genomic region 5’ of the insertion site, the 42 bp OLLAS sequence, and then a minimum of 35 bp homology arms to the genomic region 3’ of the insertion site (Supplemental Table 4). Appropriate silent mutations were included in the ssODN to prevent recutting of the edited sequence by the crRNA. As the OLLAS sequence contains a SacI restriction enzyme site, PCR screening to confirm *rpn-6.1::OLLAS* edits was followed by SacI restriction enzyme digest and agarose gel electrophoresis.

### RNA interference (RNAi) treatment

RNAi treatments were done via RNAi feeding as previously described (Timmons, Court and Fire, 2001; Allen, Nesmith and Golden, 2014; Boateng *et al.*, 2017). RNAi clones were obtained from either the Ahringer RNAi library (*rpn-1, rpn-10, rpn-13, dss-1, rpt-1, rpt-3, rpt-6, pbs-2*, and *pbs-4*) or Open Biosystems ORF-RNAi library (Huntsville, AL) (*smd-1, wee-1.3, cdk-1, rpn-2, rpn-3, rpn-6.1, rpn-7, rpn-9, rpn-11, rpn-12, rpt-2, rpt-4*, and *rpt-5*). RNAi clones for *rpn-8* and *rpn-*5 were generated in the lab (see below for details). All RNAi clones were freshly transformed into *E. coli* strain HT115 cells before usage. Either the L4440 empty vector or *smd-1(RNAi)* were used as a control RNAi condition for all RNAi treatments. *smd-1(RNAi)* was utilized because it activates the RNAi response yet has no reported reproductive phenotype in a wild-type genetic background. RNAi co-depletions were performed by measuring the optical density at 600nm wavelength of the RNAi overnight culture for each construct and then mixing the cultures in 1:1 ratio. We performed RNAi knockdown of the genes of interest by feeding the worms for a total of either 24 hours at 24°C starting from L4 stage (Figures 1–2, 6, 8 and Supplemental Figures 1-2) or 48 hours, from larval stage 4 (L4) to day 2 adult at 20°C (Figures 3–5 and Supplemental Figures 3-5) as indicated.

**Figure 2.**
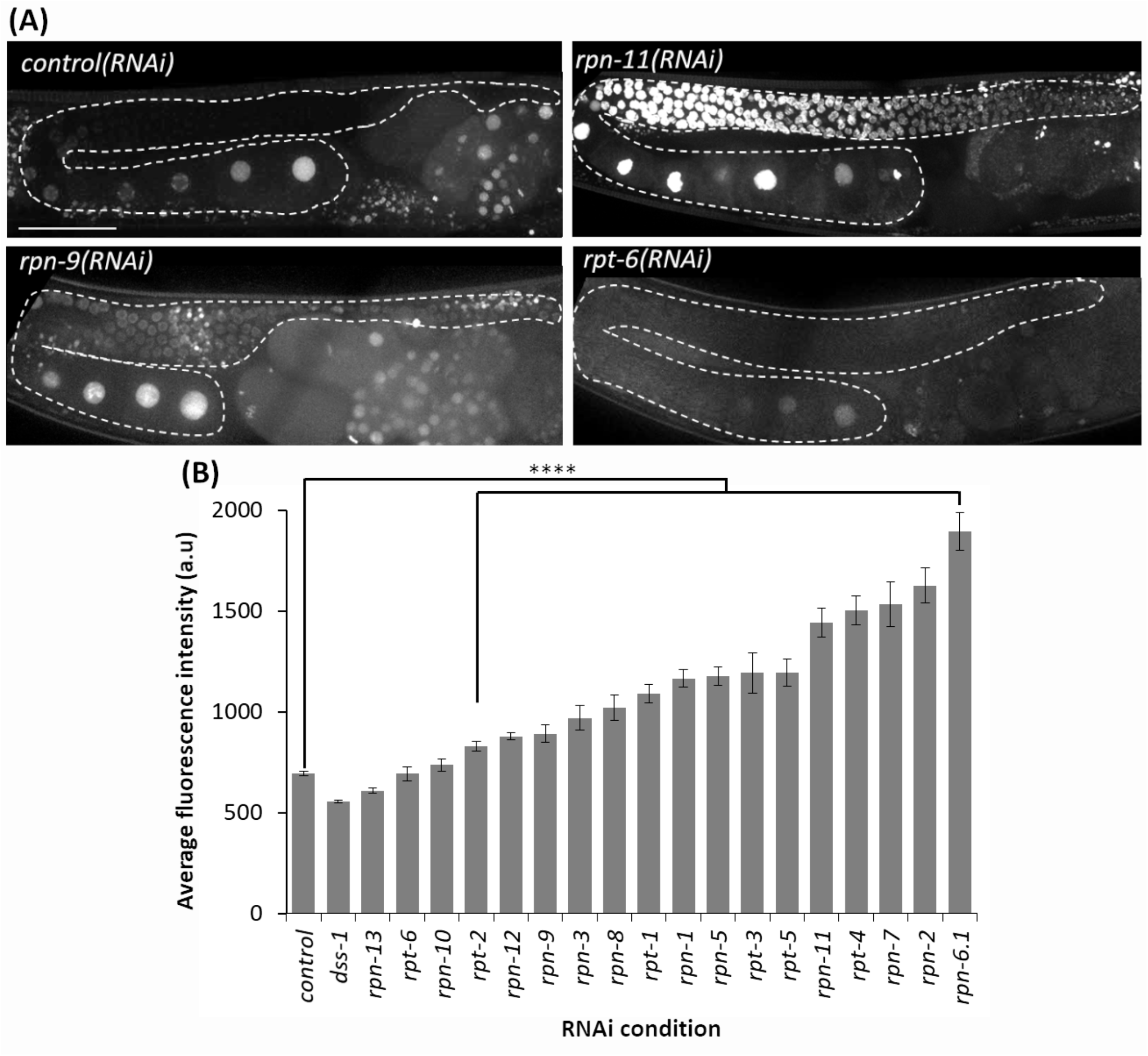
Depletion of most 19S RP subunits severely decreases proteolytic activity. (A) Germ line images of Ub(G76V)::GFP::H2B animals treated with the indicated RNAi. Representatives images of normal germline proteolytic activity [*control(RNAi)* and *rpt-6(RNAi)*], severe dysfunction of proteolytic activity [*rpn-11(RNAi)*], and moderate dysfunction of proteolytic activity [*rpn-9(RNAi)*]. A gonad arm is outlined with white dashed lines. (B) Average fluorescence intensity of Ub(G76V)::GFP::H2B germ lines treated with either RNAi against a control (n=122) or any of the various 19 subunits of the 19S RP (n=10-52). Fluorescence intensity (a.u) was measured in the region outlined with the white dashed lines as indicated in (A). All images taken at the same laser intensity and PMT gain, and then the same post-image modifications made to each image. **** represents p-values < 0.0001 compared to *control(RNAi)* condition. Error bars represent SEM. Scale bar, 50µm.

**Figure 3:**
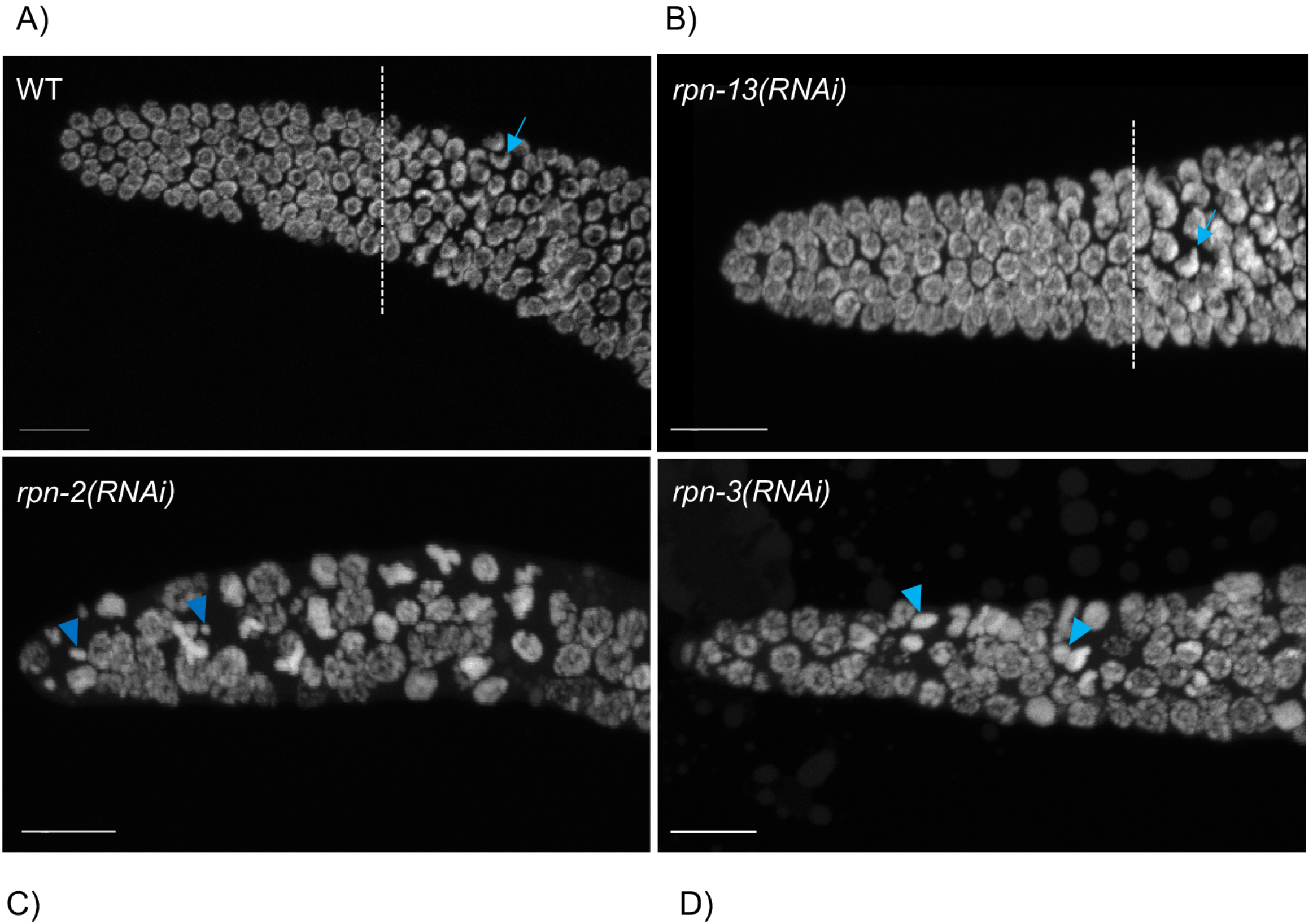
Defects in the mitotic germ line result from 19S RP subunit knockdown. Representative images of the distal tip of the *C. elegans* germ line visualized with DAPI. (A) Wild type N2 controls. (B) *rpn-13(RNAi)* resulted in no cell cycle defects, presenting mitotic tips comparable to WT worms. Both wild type and *rpn-13(RNAi)* germ lines exhibited obvious transition zones (white dash line indicates start of transition zone) with characteristic crescent shape nuclei (blue arrow), (C, D) Worms treated with *rpn-2(RNAi)* or *rpn-3(RNAi)* presented an increased number of cells in M phase and the presence of small or fragmented nuclei (blue arrowheads). Both also had shorter mitotic tips with no clear transition zone. Images show max projections of Z stacks halfway through each gonad. Scale bar, 10 µm.

**Figure 4:**
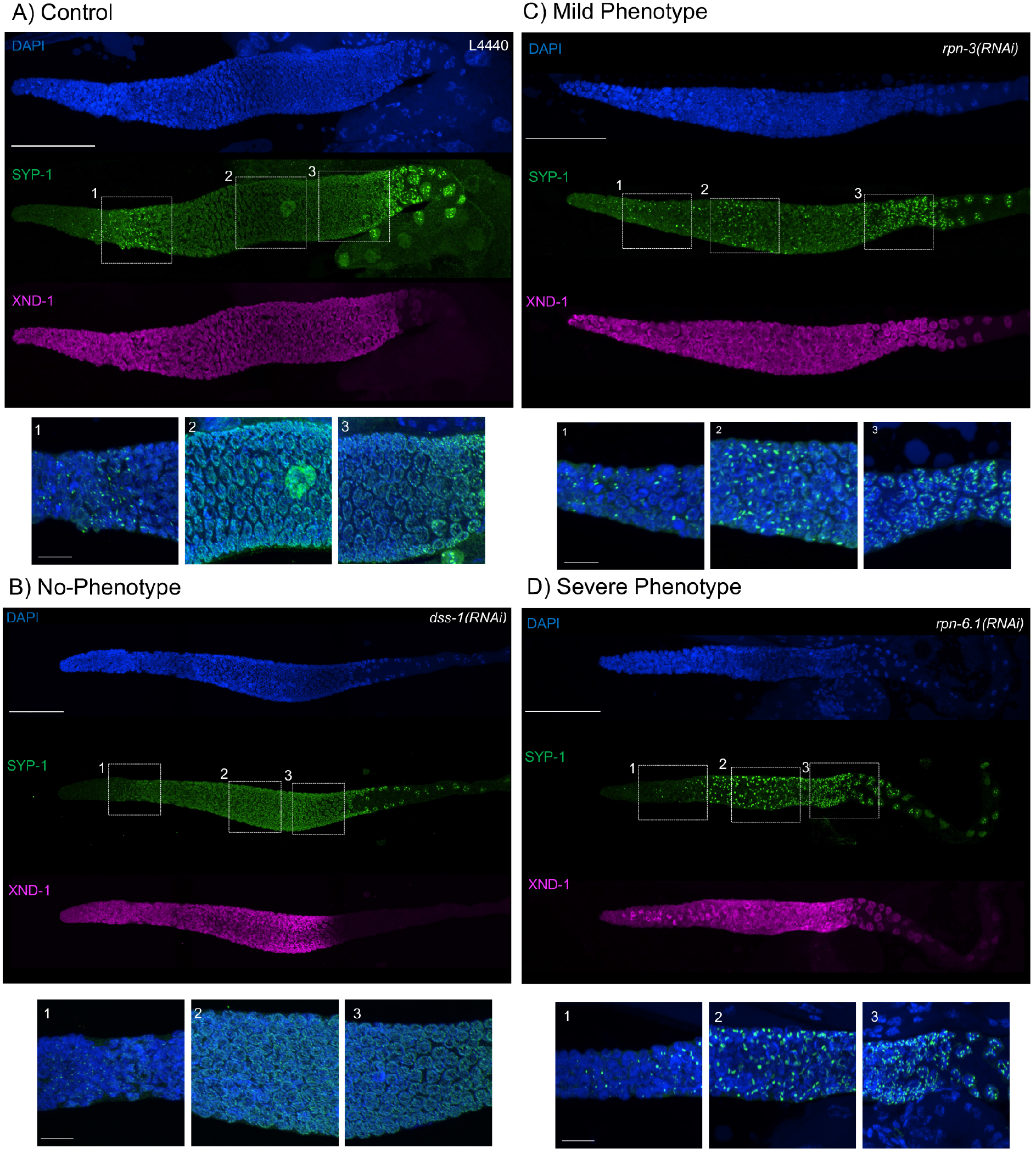
Synaptonemal complex defects are observed upon knockdown of 19S proteasome subunits. Representative images of germ lines visualized with anti-SYP-1 to mark the synaptonemal complex (green), anti-XND-1 (purple), and DAPI to mark DNA (blue). (A) Control, empty vector, shows the expected formation of a few SC polycomplexes (PCs) in TZ. (B) Mild-phenotype: extended region of PCs reaching early pachytene, with an abundant number of nuclei with fully polymerized SC in mid-pachytene. Premature polarization is also observed. (C) Severe phenotype: extended region of PCs into mid-pachytene, with almost all nuclei having at least one PC and no polymerization of SYP-1. Premature polarization of SYP-1 was present at late pachytene. (D) No phenotype: full polymerization of SYP-1 throughout pachytene stage and correct timing of polarization to the short arm of the chromosome at diplotene comparable to control. Whole gonad scale bar, 50 µm. Zoom in boxes correspond to: (1) Transition Zone, (2) Early-Mid Pachytene, (3) Late Pachytene, scale bar,10 µm

**Figure 5:**
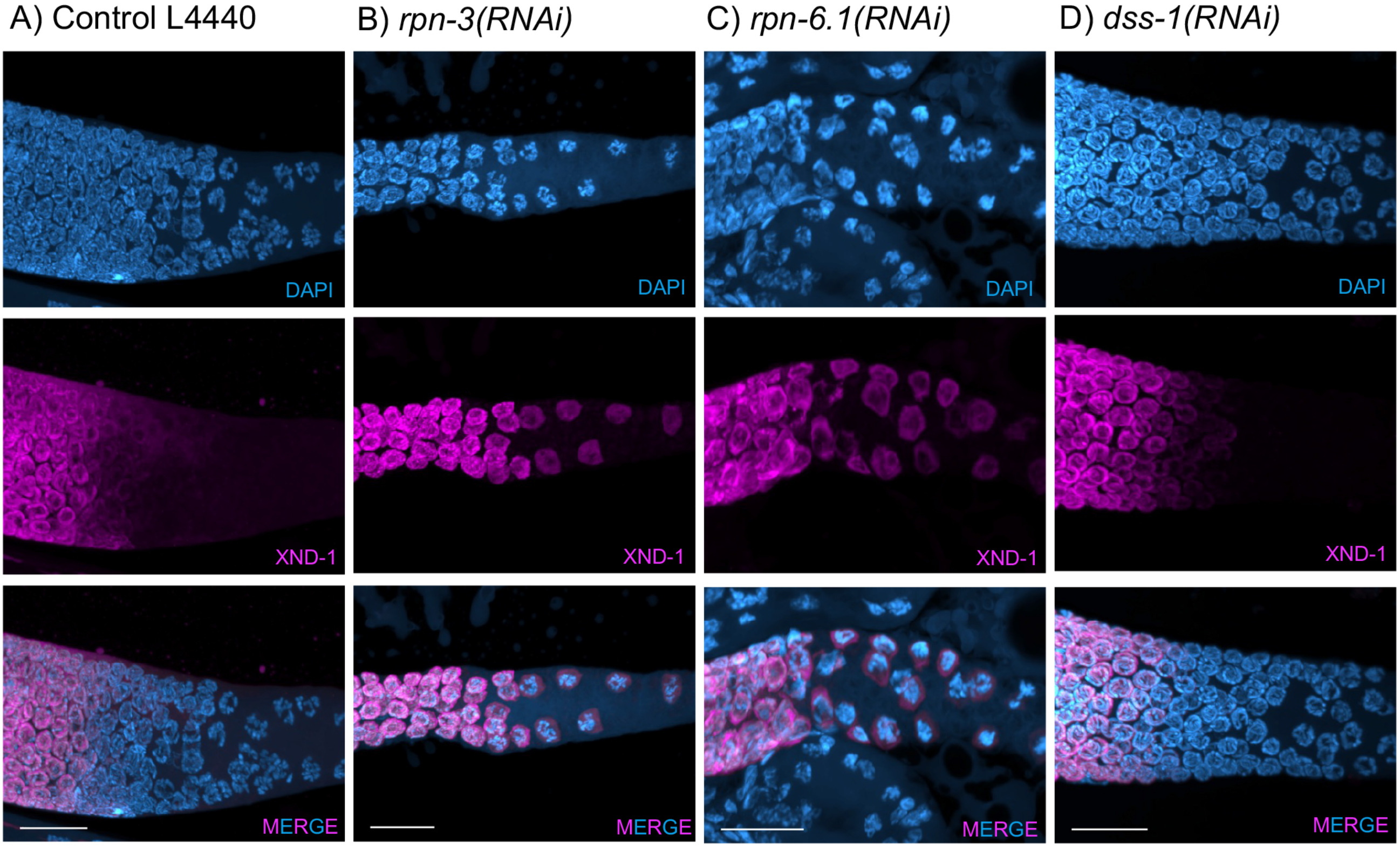
XND-1 turnover is affected by knockdown of a subset of 19S RP non-ATPase subunits. Representative images showing defects in XND-1 turnover after depletion of a specific group of non-ATPase proteasome subunits. Anti-XND-1 (magenta); DAPI stained DNA (cyan). (A) Vector control. (B) *rpn-3(RNAi)* and (C) *rpn-6.1(RNAi)* are examples of two subunits whose knockdown causes persistence of high levels of nucleoplasmic XND-1 in late pachytene nuclei. (D) *dss-1(RNAi)* is representative of the class of subunits who depletion does not affect XND-1.

**Figure 6.**
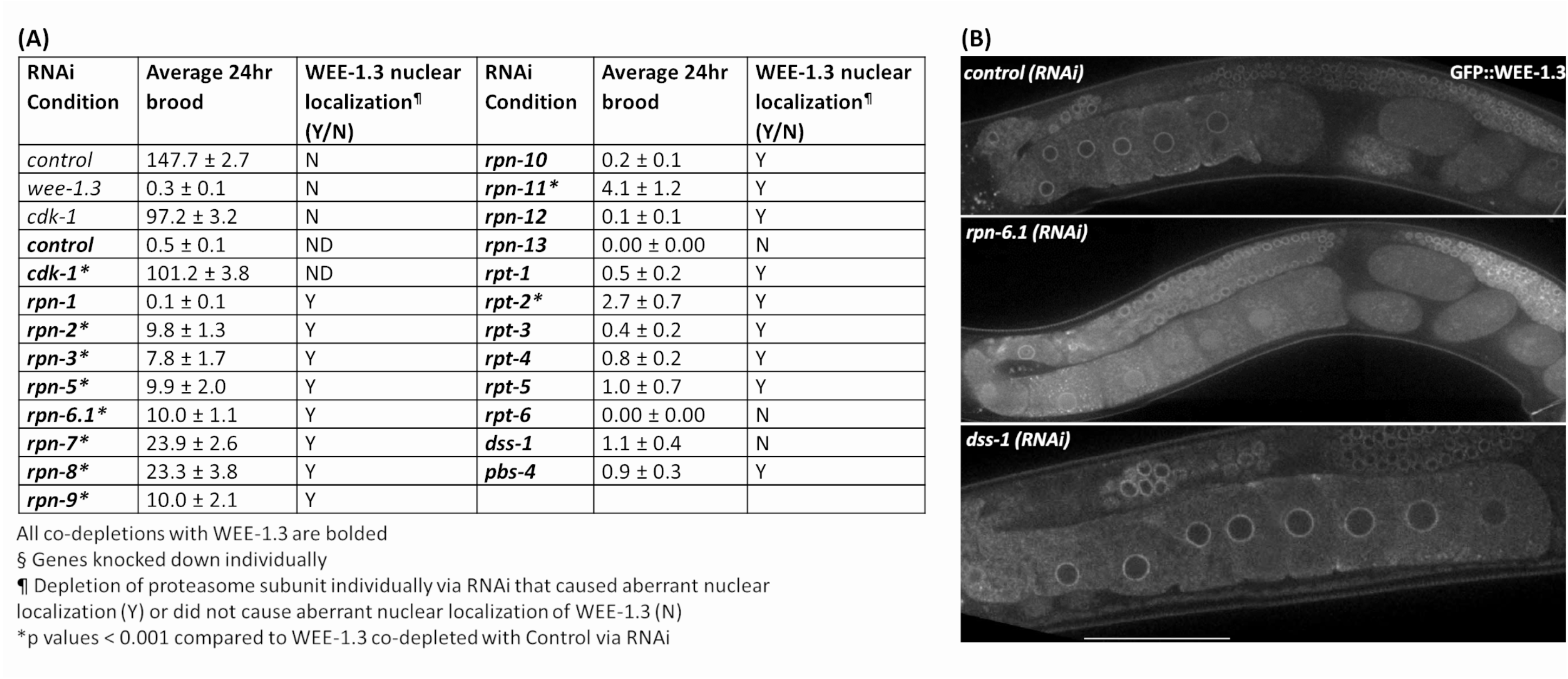
WEE-1.3 function and localization are altered by depletion of specific proteasome subunits. (A) Average 24 hr brood and WEE-1.3 nuclear localization status of hermaphrodites treated with either *control(RNAi), wee-1.3(RNAi), cdk-1(RNAi)* individually (bolded) or co-depleted with WEE-1.3, or 19S RP subunits co-depleted with WEE-1.3 via RNAi. All co-depletion conditions were compared to WEE-1.3 co-depleted with the control RNAi condition. * represents p values <0.001, Y (yes) or N (No) represents whether or not aberrant nuclear localization of WEE-1.3 occur when control or proteasome subunits depleted individually. (B) Live imaging of germ lines from strain WDC2 *wee-1.3(ana2[gfp::wee-1.3])* treated with either *control(RNAi), rpn-6.1(RNAi)* or *dss-1(RNAi)*. All images were taken at the same laser intensity and PMT gain. Scale bar, 100µm.

**Figure 7.**
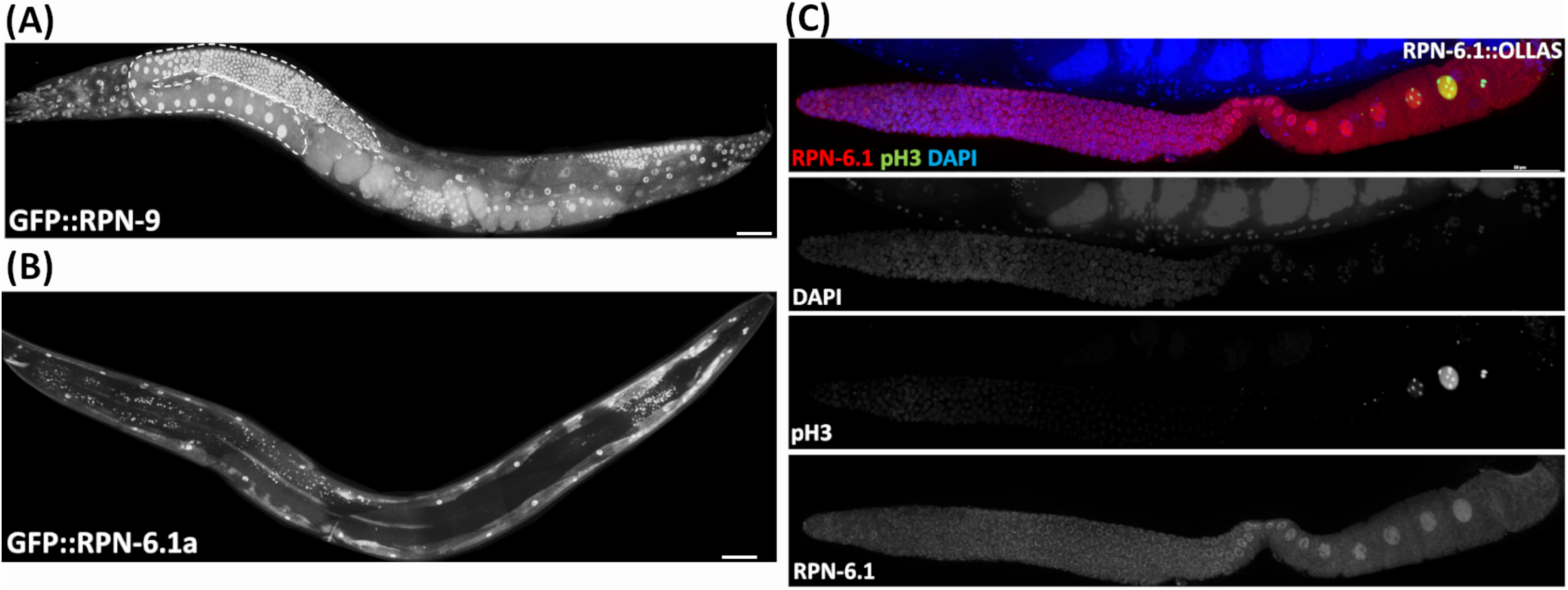
The two RPN-6.1 isoforms exhibit different spatial localization. Live imaging of hermaphrodites expressing endogenously GFP-tagged (A) RPN-9 and (B) RPN-6. Strains are WDC5 *rpn-9(ana5 [gfp::rpn-9])* and WDC3 *rpn-6.1a*(*ana3[gfp::rpn-6.1a])*. C) Immunofluorescence image of *rpn-6.1(ana12[rpn-6.1::ollas])* strain dissected germ line co-stained with anti-OLLAS (red), anti-pH3 (green, condensed chromatin) and DAPI for DNA (blue). Bright nuclear and relatively dim cytoplasmic RPN-6.1b expression shown throughout germ line. Scale bar, 50µm.

**Figure 8.**
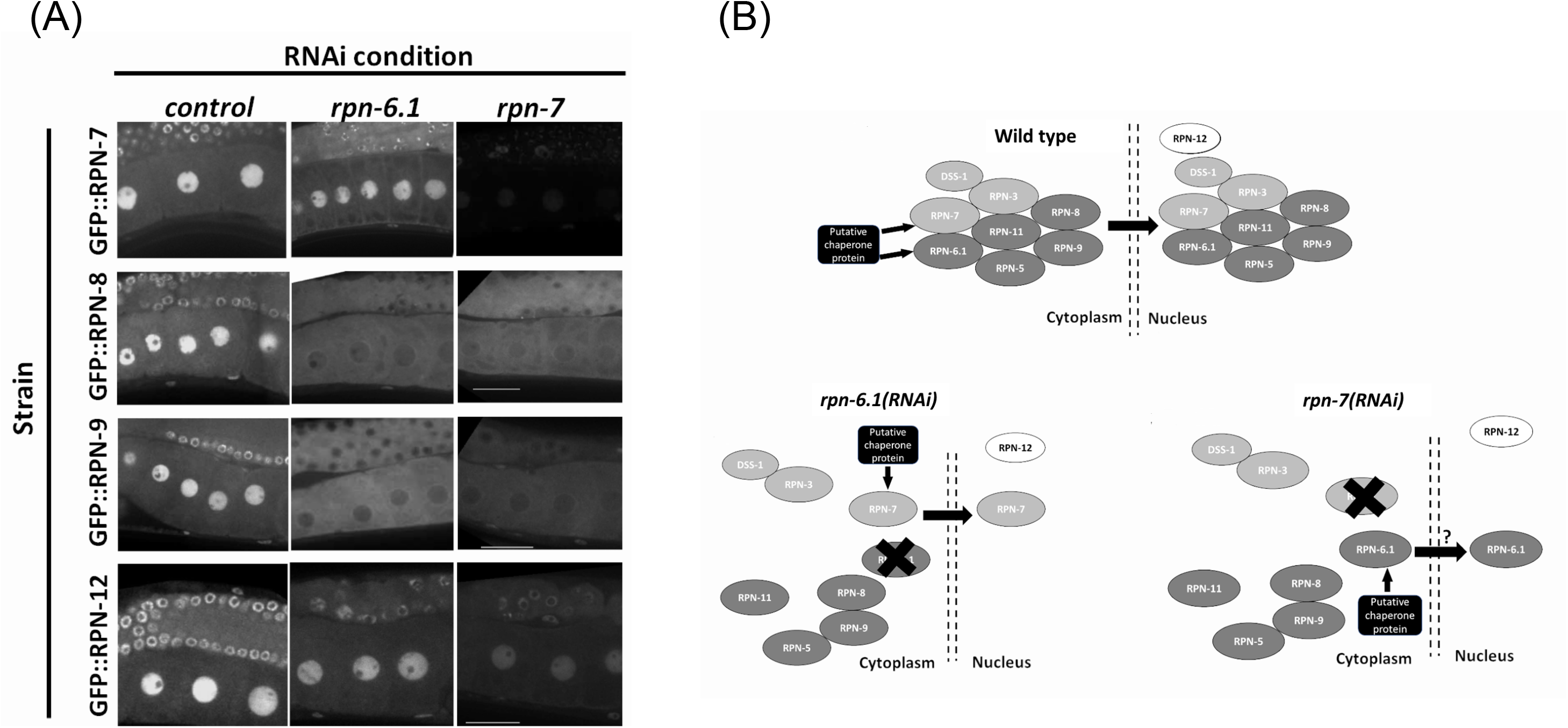
RPN-6.1 and RPN-7 are required for the nuclear localization of RPN-8 and RPN-9. (A) Live imaging of hermaphrodite oocytes from endogenously GFP tagged strains *rpn-7(ana1[gfp::rpn7])*, *rpn-8(ana4[gfp::rpn-8])*, *rpn-9(ana5[gfp::rpn-9])* and *rpn-12(ana6[gfp::rpn-12])* treated with either *control(RNAi), rpn-6.1(RNAi)* or *rpn-7(RNAi)* (n = 15 - 42). Scale bar represents 25µm. (B) Model for role of RPN-6.1 and RPN-7 in nuclear localization of 19S RP lid combining existing information on eukaryotic proteasome assembly model (Budenholzer *et al.*, 2017; Bai *et al.*, 2019).

### RNAi clone generation

RNAi feeding clones for *rpn-5* and *rpn-8* were generated by TA cloning a PCR product containing a genomic sequence of the appropriate gene into the MCS of pL4440 RNAi feeding vector. To generate clones, a 1143bp region of *rpn-5* and 504bp region of *rpn-8* was PCR amplified using MyTaq™ DNA Polymerase (Bioline Cat. No. 21105). The following primers were used: for *rpn-5*, forward oAKA277 5’-aatggctatcgcaaagatgg-3’ and oAKA278 reverse 5’-gtcagtttgtgcacgttgct-3’; and for *rpn-8*, forward oAKA392 5’-gcgtttctcactgttatgtcg −3’ and reverse oAKA393 5’-ccatgtcgaggaaccatgta-3’. In brief, the vector was linearized with EcoRV, gel-extracted (Bioline Cat. No. BIO-52059), T-tailed, desalted with a DNA Clean Concentrator kit (Zymo Research Cat. No. D4004), and then ligated with either of the previously mentioned PCR product using Quick-Stick ligase (Bioline Cat. No. BIO-27027). Newly generated RNAi clones were transformed into HT115 cells and sequenced using the M13 forward universal primer to confirm successful cloning (Eurofins Genomics).

### Fertility assays

24-hour total brood assays on RNAi-treated worms were performed using the previously published protocol with a minimum of 3 independent trials (Boateng *et al.*, 2017). Statistical analyses were performed in Microsoft Excel using the Student *T*-test to find significant differences between the average 24-hour brood of control and experimental RNAi conditions. Standard error of the mean (SEM) was calculated by dividing the standard deviation by the square root of the sample size.

### Live Imaging

All fluorescent strains were treated with appropriate RNAi condition at 24°C for 24hrs before imaging. 10µl of anesthetic (0.1% tricane and 0.01% tetramisole in 1X M9 buffer) was added to a 3% agar pad on a slide and 10-15 live worms were transferred to the drop of anesthetic. A glass coverslip was slowly lowered to cover the samples and the coverslip edges were sealed with nail polish and allowed to dry before imaging. Images were obtained on a Nikon Ti-E-PFS inverted spinning-disk confocal microscope using a 60x 1.4NA Plan Apo Lambda objective. The microscope consists of a Yokowaga CSU-X1 spinning disk unit, a self-contained 4-line laser module (excitation at 405, 488, 561, and 640nm), and an Andor iXon 897 EMCDD camera. Fluorescence intensities were quantified and image editing done using NIS-elements software.

### Immunofluorescence of Proximal Germline

The tube staining method was performed on dissected gonads fixed in 3% paraformaldehyde and methanol (Chen and Arur, 2017). The samples are washed using 1X PBST (0.1% tween), blocked with 30% NGS and incubated with primary antibodies at 4ºC overnight. Appropriate secondary antibodies were added and incubated at room temperature for 1-2 hours followed by three washes with 1X PBST with DAPI included in the final wash and samples were mounted on a 3% agar pad with Vectashield mounting medium. The primary antibodies used in this study are: Rat monoclonal OLLAS epitope tag antibody (1:200, Novus Biologicals, Cat. No. NBP1-06713) and Rabbit anti-phospho-Histone H3 (Ser10) antibody (1:200, EMD Millipore Cat. No. 06-570). Secondary antibodies were goat-anti-rat Alexa Fluor 568nm and goat-anti-rabbit Alexa Fluor 488 (1:1000, Invitrogen).

### Immunofluorescence of Synapsis Phenotypes in Distal Germline

For the study of synapsis, germ lines from N2 worms exposed to 48 hours RNAi by feeding, were dissected in 1x Sperm Salt Buffer (50 mM PIPES pH 7.0, 25 mM KCl, 1 mM MgSO_4_, 45 mM NaCl, 2 mM CaCl_2_), followed by permeabilization with 2% Triton and then fixed in the same buffer containing 2% paraformaldehyde for 5 min. Slides were placed on a frosted metal plate on dry ice before removing the coverslip and then placed in 4°C absolute ethanol for 1 min. Slides were then washed three times for 10 min each in PBST (1x PBS, 0.1% Tween) plus 0.1% BSA and incubated overnight at 4ºC with the primary antibodies diluted in PBST. Following three washes of 10 min each in PBST plus 0.1% BSA, slides were incubated in the dark at room temperature for 2 hours with secondary antibodies diluted in PBST. Following three 10 min washes with PBST, slides were counterstained with DAPI in the second wash and mounted using Prolong Gold antifade reagent with DAPI (Invitrogen). The primary antibodies used in this study are: Chicken anti-SYP-1 (1:1000, courtesy of Dr. Enrique Martinez-Perez) and Guinea Pig anti-XND-1 (1:2000) (Wagner *et al.*, 2010; Silva *et al.*, 2014). XND-1, a chromatin factor responsible for the global distribution of crossovers in *C. elegans*, was used as a control of the staining protocol allowing us also to identify the late pachytene stage in the germline. Secondary antibodies were goat-anti-chicken Alexa Fluor 488nm (1:2000, Invitrogen) and goat-anti-guinea pig Alexa Fluor 633nm (1:2000, Invitrogen).

## Results

### Differential roles of 19S RP subunits in *C. elegans* reproduction and larval growth observed when downregulated individually via RNAi

We wanted to compare the effects of downregulation of each of the 19S RP lid and base subunits in *C. elegans* hermaphrodites. As expected RNAi knockdown of proteasome subunits led to significant brood size reductions compared to control RNAi (Figure 1C, p value < 0.01). Whereas the majority of 19S base subunit-knockdown animals had fewer than 6 offspring (<0.4% of control), *rpt-6(RNAi)* and *rpn-13(RNAi)* animals produced substantial numbers of eggs (~25% and ~63% of controls, Figure 1C) many of which hatched (Figure 1D). By contrast, knockdown of only half of the proteasome lid subunits severely reduced broods (<10 eggs); the remainder gave brood sizes 30-80% the size of controls (Figure 1C). Of those with substantial numbers of eggs, *rpn-5* severely reduced hatching, leading to few to no viable offspring (Figure 1D). These results replicate the findings of Takahashi et al (Takahashi *et al.*, 2002). In some instances, such as *rpt-6(RNAi) and rpn-9(RNAi)*, the hatched embryos develop into larvae but exhibit severe developmental defects, such as L1-L2 developmental arrest and a protruded vulva phenotype, respectively (data not shown). This data, combined with previously published data, suggests while most of the lid and base subunits of 19S RP of the 26S proteasome play essential roles during *C. elegans* hermaphrodite reproduction, individual 19S RP subunits may play differential roles in this process.

### Downregulation of most, but not all, 19S RP subunits causes dysfunction of the proteolytic activity of the proteasome

*In vivo* fluorescent reporter systems have been developed to qualitatively assess the proteolytic activity of the 26S proteasome in whole animals and in specific tissues under various conditions (Pispa, Matilainen and Holmberg, 2020). This technique takes advantage of a translational fusion of a mutated, non-hydrolysable ubiquitin moiety to a fluorescent reporter, thereby subjecting the fluorescent protein to continuous proteasomal degradation (Dantuma *et al.*, 2000; Hamer, Matilainen and Holmberg, 2010; Liu *et al.*, 2012). Here, we use the published IT1187 strain with a mutated ubiquitin fused to a GFP-tagged histone protein and driven by a germline specific promoter (*pie-1_pro_*::Ub(G76V)::GFP::H2B::*drp-1* 3’UTR) (Kumar and Subramaniam, 2018). GFP can thus be used as an indicator of germline proteolytic activity upon RNAi depletion of specific 19S RP subunits (Fernando, Elliot and Allen, 2020). If the proteolytic activity of the proteasome is normal, the non-hydrolysable mutated ubiquitin will target the GFP::H2B for continuous proteasomal degradation leading to dim or no GFP signal in the hermaphrodite germ line. Dysfunction of the proteolytic activity of the 26S proteasome leads to accumulation of Ub(G76V)::GFP::H2B resulting in bright GFP.

RNAi depletion of all of the lid subunits except *rpn-10*, *rpn-13*, *dss-1/rpn-15*, and *rpt-6* resulted in bright, nuclear, germline fluorescence of the Ub(G76V)::GFP reporter compared to control RNAi-treated germ lines (Figure 2A and Supplemental Figure 1). To compare proteolytic activity of these components, we quantified the GFP intensity in germ lines depleted of specific 19S RP subunits and imaged them under the same microscopy conditions (Figure 2B). This confirmed our qualitative observations that RNAi depletion of lid subunits does not uniformly impact germline proteolytic activity. For example, depletion of *rpt-2*, *rpn-9* or *rpn-12* resulted in only a modest increase in GFP fluorescence whereas RNAi of *rpn-2, rpn-7*, and *rpn-6.1* exhibited the greatest increase in fluorescence (Figure 2B). One trivial explanation for these differences in fluorescence and phenotypes are differential sensitivity of the proteasome genes to RNAi perturbation. We do not favor this explanation at least for *rpn-9* and *rpn-12*: our fluorescent reporters (described below) allowed us to ascertain that subunit expression can be effectively inhibited even for those subunits where we observe little to no phenotypic changes (Supplemental Figure 2). Therefore, we speculate that specific 19S RP proteasome subunits may contribute uniquely to the proteolytic activity in the germ line.

### Downregulation of specific 19S RP subunits causes cell cycle defects in the adult germ line

The ubiquitin proteasome system plays a central role in cell cycle regulation (reviewed in (Zou and Lin, 2021)). In the *C. elegans* germ line, the mitotic cells reside in the distal tip, or proliferative zone (PZ), and provide the pool of cells that enter meiosis as they move proximally (Figure 1B). Under normal growth conditions on day one of adulthood, ~2.5% of cells have been reported to be in M phase based on staining with phospho-histone H3 (Kocsisova, Kornfeld and Schedl, 2019). Accordingly, under control RNAi conditions, we observed only rare metaphase or anaphase figures in the mitotic zone (Figure 3). By contrast, upon RNAi knockdown of most of the lid subunits (*rpn-3*, *rpn-5*, *rpn-6.1*, *rpn-7*, *rpn-8, rpn-9*, or *rpn-11*) and the base subunits *rpn-1* and *rpn-2*, we observed increased numbers of cells at metaphase or anaphase (Figure 3, Table 1, and Supplemental Figures 3, 4). We also observed severe defects in the PZ nuclei that are never seen in wild type: very small nuclei, fragmented nuclei, and chromosome fragments (Figure 3, arrowheads). Overall, these RNAi exposures led to shorter PZs with heterodisperse nuclear sizes and shapes compared to the orderly and uniform mitotic regions of controls. These phenotypes were also accompanied by a change in nuclear morphology at meiotic entry. In wild-type and control RNAi-exposed animals, the transition zone (TZ) nuclei (corresponding to leptotene/zygotene stages of meiosis) have a distinctive crescent shape (Hillers *et al.*, 2015). After 48h of exposure to proteasome RNAi, the TZ nuclei were difficult to distinguish from the anaphase-like chromosomes in the mitotic region (Crittenden *et al.*, 2006; Hubbard, 2007) (Figure 3). In contrast to the profound proliferative defects described above, RNAi knockdown of the non-ATPase subunits *rpn-10*, *rpn-12*, *rpn-13* and *dss-1/rpn-15* did not alter PZ nuclear size or morphology and they appeared indistinguishable from control worms in this region (Supplemental Figure 5 and data not shown).

### Downregulation of specific 19S RP subunits compromises both SC assembly and SC reorganization in late pachytene

Previous work from our group and others has shown that a structurally compromised proteasome core complex results in severe defects both in synaptonemal complex (SC) assembly and in premature reorganization of the SC in late pachytene (Ahuja *et al.*, 2017; Prasada Rao *et al.*, 2017). Based on these results, we wanted to interrogate how these events are affected when the 19S RP subunits are knocked down. In TZ nuclei, the SC central region proteins self-aggregate forming polycomplexes (PCs) (Goldstein, 1986). These PCs can be seen as bright foci using immunofluorescence or live imaging of fluorescently-tagged SC proteins (Figure 4) (Rog, Köhler and Dernburg, 2017). In wild type, PCs can be seen only in ~one to four nuclei because they disappear as the SC proteins polymerize along chromosomes to form the SC (Figure 4A) (Rog, Köhler and Dernburg, 2017) The PC region is extended if the SC cannot polymerize, for example due to defects in SC regulatory proteins, among others (Couteau and Zetka, 2005; Martinez-Perez and Villeneuve, 2005). Similar to what we observed with knockdown of the 20S subunit, RNAi knockdown of *rpn-1*, *rpn-2*, *rpn-3*, *rpn-5*, *rpn-6.1*, *rpn-7*, *rpn-8* or *rpn-11* resulted in an extended region of SYP-1 PCs (Figure 4C, D, Supplemental Figures 3, 4) (Ahuja *et al.*, 2017). As shown in Figure 4, both the number of nuclei that have PCs and the size of the PCs was increased in knockdown animals after 48hr of proteasome RNAi compared to control RNAi (Figure 4C, D). In the nuclei where PC persist, little to no SC is seen on chromosomes. In the most severe germ lines, PCs can be seen into mid-pachytene, well into the region that would normally be fully synapsed (compare Figure 4D vs 4A). In contrast to the robust phenotypes described above, the knockdown of the remainder of the non-ATPase subunits (*rpn-9*, *rpn-10*, *rpn-12*, *rpn-13* or *dss-1*) had no obvious effect on SC assembly or on PC size, number, or persistence (Figure 4B, Supplemental Figure 5). We note that *rpn-9* is distinct in having effects on mitotic proliferation but not on PC turnover/SC assembly.

In late pachytene, remodeling of SC occurs to facilitate bivalent formation: SYP proteins are removed from the long arm of the chromosome (relative to the crossover) and are retained and enriched on the short arm (MacQueen *et al.*, 2002; Colaiácovo *et al.*, 2003). The remodeling first becomes apparent in late pachytene nuclei by polarization of SC subunit into bright and dim patches seen by immunofluorescence (MacQueen *et al.*, 2002; Colaiácovo *et al.*, 2003). In the proteasome 20S knockdown, we observed premature polarization of SYP with patches appearing more distally than in the wild-type controls (Ahuja *et al.*, 2017). Upon 19S RP subunit RNAi, we saw complete congruence between subunits that showed early PCs and those that presented with premature polarization (Figure 4C, D, Supplemental Figures 3, 4). In the most severe RNAi exposures, the polarization began in the mid-pachytene region (Figure 4D, Supplemental Figure 3). Similarly, those genes whose knockdown did not result in accumulation of PCs also did not show the premature polarization of the SC (Figure 4B, Supplemental Figure 5).

### Nuclear XND-1 levels are regulated by the proteasome

In addition to the effects previously described for proteasome inhibition in the meiotic region of the germ line, we also observed that the proteasome is required for the proper down-regulation of XND-1 (X non-disjunction factor 1) protein in late pachytene (Figure 5). XND-1 is a chromatin factor, responsible for the global distribution of meiotic crossovers in *C. elegans* (Wagner *et al.*, 2010). In wild type, XND-1 protein is localized on autosomes from the mitotic tip of the germ line until late pachytene (Wagner *et al.*, 2010). At that time, XND-1 appears to dissociate from chromosomes and the nuclear XND-1 signal diminishes. In cellularized oocytes, prior to ovulation, the predominant pool of XND-1 protein is cytoplasmic where it remains until it is ultimately segregated into the developing germ cells of the embryo (Mainpal, Nance and Yanowitz, 2015). In contrast to wild-type and control RNAi-exposed animals, we observed that knockdown of *rpn-1*, *rpn-2*, *rpn-3*, *rpn-5*, *rpn-6.1*, *rpn-7*, *rpn-8* or *rpn-11*, the same subunits that altered the SC polymerization and restructuring, also led to defects in XND-1 turnover. In the late pachytene nuclei of these RNAi-exposed animals, XND-1 levels remained high and nucleoplasmic (Figure 5). Thus, we infer that these subunits are not required for the chromosomal association of XND-1 *per se*, but rather are responsible for the turnover and/or export of the non-chromosomally associated XND-1 pool.

### Downregulation of specific 19S RP subunits suppresses *wee-1.3(RNAi)* infertility and alters WEE-1.3 localization in oocytes

*C. elegans* oocytes, like oocytes of most sexually reproducing organisms, undergo meiotic arrest (Burrows *et al.*, 2006; Inoue *et al.*, 2006; Ruiz, Vilar and Nebreda, 2010). Oocyte meiotic arrest in *C. elegans* hermaphrodites is maintained by an inhibitory kinase WEE-1.3 phosphorylating the CDK-1 component of maturation promoting factor (MPF) and thus inactivating MPF (Lamitina and L’Hernault, 2002; Burrows *et al.*, 2006; Allen, Nesmith and Golden, 2014). Depletion of WEE-1.3 in *C. elegans* causes precocious oocyte maturation resulting in infertility (Burrows *et al.*, 2006). A large RNAi suppressor screen identified 44 suppressors that when co-depleted with WEE-1.3 suppressed the infertility defect (Allen, Nesmith and Golden, 2014). Five of the suppressor genes were subunits of the 19S RP. However not all of the 19S RP subunits were included, or identified as positives, in the aforementioned screen (Allen, Nesmith and Golden, 2014). Therefore, we systematically screened each of the 19S RP subunits to determine if there are additional subunits whose depletion suppresses *wee-1.3(RNAi)* induced infertility.

Hermaphrodites fed *wee-1.3(RNAi*) are infertile, averaging less than one egg per adult hermaphrodite (Figure 6). In the absence of CDK-1, WEE-1.3 is dispensable. Accordingly, *cdk-1(RNAi)* suppresses *wee-1.3(RNAi)* infertility and therefore serves as a positive control in these studies (Figure 6A) (Burrows *et al.*, 2006). Significant increases in brood sizes were seen when WEE-1.3 was co-depleted with 8 out of 13 of the 19S lid subunits, but only seen with co-depletion of one of the 19S base subunits, RPT-2 (Figure 6A). Depletion of the remaining 5 base units were unable to suppress, similar to the negative control co-depleted with WEE-1.3 (Figure 6A).

WEE-1.3 is mainly localized to the perinuclear region, but also can be seen in the cytoplasm and ER (Allen, Nesmith and Golden, 2014). Depletion of most 19S RP subunits in an endogenously GFP tagged WEE-1.3 strain [WDC2 – *gfp::wee-1.3(ana2)*] caused aberrant nuclear accumulation of WEE-1.3 (Figure 6B). RNAi of four of the 19S RP subunits that failed to suppress *wee-1.3(RNAi)* sterility, RPN-10, RPN-13, DSS-1/RPN-15 and RPT-6, also showed no change in GFP::WEE-1.3 localization (Figure 6B, Table 2, and data not shown). However, since we previously reported that *rpn-10(ana7)*, a genetic null, results in nuclear accumulation of GFP::WEE-1.3 in oocytes, it is possible that our RNAi depletions of RPN-13, DSS-1 or RPT-6 did not give sufficient knockdown to elicit an alteration in perinuclear WEE-1.3 localization (Fernando, Elliot and Allen, 2020). However, our previous study also reported that chemical inhibition of the proteolytic activity of the proteasome with Bortezomib neither suppressed *wee-1.3(RNAi)* infertility nor induced nuclear accumulation of WEE-1.3 (Fernando, Elliot and Allen, 2020). Therefore, we favor the conclusion that a fully intact 19S RP is required for the proper localization of WEE-1.3 in oocytes and that this role is independent of the proteasome’s role in proteolysis.

### Ubiquitous somatic and germline expression of 19S RP lid subunits RPN-7, RPN-8, and RPN-9

The transparency of *C. elegans* makes it an excellent model to conduct live imaging of fluorescently tagged proteins and is useful to study highly dynamic protein complexes such as the 26S proteasome. To better understand the spatiotemporal expression of 19S RP subunits *in vivo* and ultimately to perform future biochemical analyses, we set out to endogenously tag each of the 19S RP subunits with GFP or OLLAS. We previously reported that an endogenous GFP::RPN-12 strain exhibits somatic and germline expression (Fernando, Elliot and Allen, 2020). N-terminal GFP fusions with RPN-7, RPN-8, or RPN-9 showed ubiquitous expression in both the nuclei and cytoplasm of germline and somatic cells, including developing oocytes (Figure 7A and Supplemental Figure 6). This subcellular expression matches that determined by antibody staining against subunits of the proteasome core particle in *C. elegans* and in other systems (Brooks *et al.*, 2000; Mikkonen, Haglund and Holmberg, 2017; Kumar and Subramaniam, 2018; Fernando, Elliot and Allen, 2020). Importantly, all three of these strains exhibited no effect on lifetime brood size and only a moderate reduction in lifespan when compared to wild-type control animals (data not shown).

### Expression of the 19S RP lid subunit RPN-6.1a is restricted to the body wall muscle

While the 19S RP subunits (RPN-7, −8, −9, and −12) all exist as a single protein isoform, the RPN-6.1 subunit has two protein isoforms, A and B, that differ by an extension of the N-terminus in RPN-6.1A (Supplemental Figure 7) (Wormabse, 2022). A strain endogenously tagging the N-terminus of RPN-6.1A with GFP shows nuclear and cytoplasmic GFP expression restricted to the body wall muscle cells of the animal (Figure 3B, strain WDC3 *rpn-6.1a(ana3[gfp::rpn-6.1a])*). Since an N-terminal fusion of RPN-6.1B would impact expression of RPN-6.1A, we instead attempted to infer its expression from an endogenous GFP tag to the C-terminus of RPN-6.1, which would simultaneously tag both RPN-6.1 isoforms (Supplemental Figure 7). Unfortunately, we were unable to obtain viable or fertile RPN-6.1::GFP animals, suggesting GFP interfered with the proper folding or function of RPN-6.1. Instead, we were able to create a functional gene fusion using a small epitope tag, OLLAS (WDC12 *rpn-6.1(ana12[rpn-6.1a::ollas])*). Lifespan and lifetime brood assays of the *gfp::rpn-6.1a* and *rpn-6.1::ollas* strains demonstrated that the N-terminal tag had no effect compared to wild-type control animals, while the C-terminal OLLAS tag results in a slightly reduced lifetime average brood and lifespan compared to wild-type control (data not shown).

We immunostained dissected RPN-6.1::OLLAS animals with an anti-OLLAS antibody and as predicted, we observed staining in the nuclei and cytoplasm of germ line and intestinal cells (Figure 7C). Since GFP::RPN-6.1A fluorescence was restricted to the body wall muscle, the anti-OLLAS staining that we observed in the germ line and intestine can be inferred to be due to the expression of RPN-6.1B. Interestingly, sperm did not exhibit expression of either isoform RPN-6.1A or B. We hypothesize that this may be due to the presence of a sperm-specific ortholog of *rpn-6.1*, *rpn-6.2*, that is reported as expressed in sperm (Dr. Lynn Boyd personal communication and WormBase). Additionally, neither *gfp::rpn-6.1a* nor *rpn-6.1::ollas* animals exhibit expression in the pharynx, unlike other tagged proteasomal subunits, for example *gfp::rpn-9* (data not shown). This implies that the pharynx might either have a pharyngeal-specific proteasomal subunit orthologous to RPN-6.1 or that the pharyngeal proteasome does not utilize an RPN-6.1 subunit for function.

### RPN-6.1 and RPN-7 are required for nuclear localization of the 19S RP subcomplex

Yeast and mammalian studies have shown that the 26S proteasome can assemble in either the cytoplasm or the nucleus (Satoh *et al.*, 2001; Yashiroda *et al.*, 2008; Kaneko *et al.*, 2009; Murata, Yashiroda and Tanaka, 2009; Kish-Trier and Hill, 2013; Pack *et al.*, 2014; Bai *et al.*, 2019; Wendler and Enenkel, 2019). The subunits first assemble as subcomplexes in the cytoplasm with the help of chaperones and can then be imported into the nucleus where they combine to form the mature 26S proteasome forms (Le Tallec *et al.*, 2007; Li *et al.*, 2007; Murata, Yashiroda and Tanaka, 2009; Wendler and Enenkel, 2019). In yeast, the 20S CP subcomplexes assemble by at least five proteasome assembly chaperones, PAC1-PAC4 and POMP (Le Tallec *et al.*, 2007; Bai *et al.*, 2019). Nuclear localization of these 20S CP subcomplexes use the nuclear localization sequences (NLS) of the alpha subunits (Enenkel, 2014; Budenholzer *et al.*, 2020). The 19S RP assembly occurs in several steps where the lid and base take different routes to the nucleus before joining the 20S CP to complete 26S proteasome assembly (Isono *et al.*, 2007). The base assembly in yeast requires several chaperones, Nas6, Nas2, Hsm5 and Rpn14, and its nuclear localization is known to be carried out by NLS sequences in the RPN2 and RPT2 subunits (Wendler *et al.*, 2004; Funakoshi *et al.*, 2009; Roelofs *et al.*, 2009; Enenkel, 2014; Bai *et al.*, 2019). Meanwhile the yeast lid subcomplex forms into two intermediate modules before joining to form the full lid (Bai *et al.*, 2019). The two intermediate modules consists of RPN3, RPN7, and RPN15, and of RPN6, RPN8, RPN9, and RPN11 (Isono *et al.*, 2007; Bai *et al.*, 2019). RPN6 and RPN7 then interact to form the complete lid subcomplex, before the last lid subunit, RPN12, joins the subcomplex (Tomko and Hochstrasser, 2011). The lid subunits do not possess canonical NLS sequences, therefore the nuclear localization mechanism of the lid subcomplex remains unclear.

Our previous results demonstrated a nuclear pool of many 19S RP subunits. To test if any *C. elegans* 19S subunits are necessary for the nuclear localization of lid subcomplex components, we downregulated individual 19S RP lid subunits via RNAi and asked whether localization of other 19S RP subunits was affected. RNAi depletion of either RPN-6.1 or RPN-7, but not other lid subunits, impacted the nuclear signal of GFP::RPN-8 and GFP::RPN-9 in oocytes (Figure 8A-B, Supplemental Figure 2). By contrast, these depletions did not impact GFP::RPN-7 and GFP::RPN-12 localization (Figure 8B). Together our data show that RPN-6.1 and RPN-7 are required for the nuclear localization of the 19S RP lid particle subcomplexes.

## Discussion

We propose that the *C. elegans* germ line can serve as a model to study proteasome subunit dynamics *in vivo*. Endogenous fluorescent-labeling of specific subunits showed cellular and subcellular localization of those subunits that has not been clearly reported by previous studies. Our depletion studies for each the 19S regulatory particle subunits have uncovered catalytic and structural roles for the whole proteasome, lid-specific functions, as well as evidence for moonlighting roles of specific subunits.

Individual subunits of the 19S regulatory particle (RP) of the *C. elegans* proteasome contribute to different extents to a range of germ line processes. RNAi depletion of 13 out of 19 subunits of the 19S RP (Table 2) caused very high rates of embryonic lethality in progeny of treated mothers (hatching <20%; where 12/13 were <5%). All 13 of these subunits also caused severe impairment of the proteolytic activity of the proteasome as measured with the germ line, Ub(G76V)::GFP::H2B reporter. Eight of the 13 were tested for additional germ line defects and all exhibited impaired mitotic divisions, SC defects, aberrant WEE-1 localization, and retention of XND-1 in late pachytene nuclei. This latter phenotype is particularly noteworthy because it occurs at/near the time when a) profound changes in oocyte transcription and chromatin are occurring to prepare the oocyte for embryonic development and b) a subset of nuclei is culled by apoptosis. Whether the proteasome plays a pivotal role(s) in promoting these transitions deserves further investigation. With the exception of WEE-1.3 localization, these phenotypes were also impacted by bortezomib and knockdown of one or more core proteasome subunits. Together these data support the conclusion that proteasomal activity plays critical and essential roles throughout the *C. elegans* hermaphrodite germ line to ensure proper oocyte development and ensuing embryonic viability.

The depletion of the *rpn-9* and *rpn-12* subunits moderately impaired proteolytic activity of the proteasome (Figure 2B and Supplemental Figure 1) without severely affecting brood sizes (~50% and ~66% reductions) or hatching rates (~50% and ~20% reductions, respectively). One possible explanation is that the assays reflect differential requirements for proteasome function in different cells: Ub(G76V)::GFP expression is assayed in the meiotic germ line and developing oocytes; brood sizes reflect a combination of mitotic divisions, apoptosis, and oocyte maturation; and hatching rates reflect the impacts on the laid eggs. Consistent with this interpretation, *rpn-9(RNAi)* but not *rpn-12(RNAi)* exhibited mitotic zone defects which could explain the brood size defects in the former. Alternatively, there may be regional or cell type-specific differences in the RNAi efficiency for these subunits or different sensitivities of these phenotypic readouts to proteasome impairment. A final possibility, relating specifically to *rpn-12*, is the previously proposed idea that *rpn-10* and *rpn-12* are redundant and can compensate for one another during oocyte development (Takahashi *et al.*, 2002; Shimada *et al.*, 2006; Fernando, Elliot and Allen, 2020).

One of our surprising observations is that RNAi directed against *dss-1*, *rpn-13*, *rpn-10*, and *rpt-6* had mild to no effect on many of the processes examined. While these results may indicate that the RNAi is inefficient at knocking down these subunits, we note that all four knockdowns did have a mild effect on brood size, producing 25-80% of the number of eggs as wild type, strongly suggesting the RNAi is working. Our data and previously published studies using mutant analyses have shown that RPN-10, RPN-12 and DSS-1 play significant roles in the hermaphrodite germline sex determination pathway, oogenesis, and later on during larval development and growth (Shimada *et al.*, 2006; Pispa *et al.*, 2008; Fernando, Elliot and Allen, 2020). Although 99% of the embryos hatched upon knockdown of RPN-13, most larvae presented a ruptured vulva phenotype (data not shown). These data strongly suggest that RNAi depletion of these subunits is functional. One possible model for the lack of strong phenotype is that other proteostasis mechanisms may be upregulated when these subunits are inactivated, thereby supporting development and fertility with a partially compromised proteasome. Prior studies have revealed such cross-pathway feedback mechanisms, but whether all tissues respond similarly is not clear (Li, Li and Wu, 2022).

RPN-10, RPN-13 and DSS-1 are known as ubiquitin receptors of the 26S proteasome, but there is evidence to suggest that these subunits confer substrate specificity and do not function as global receptors of polyubiquitinated proteasome substrates (Shimada *et al.*, 2006; Paraskevopoulos *et al.*, 2014). In mammalian cells, RPN10 can compensate for loss of RPN13, and vice versa, presumably because of their shared role in ubiquitin-binding (Hamazaki, Hirayama and Murata, 2015). It would be interesting to test whether similar compensation happens in the worm. RPN-1 is the only other 19S RP subunit thought to have ubiquitin-binding activity. Since loss of RPN-1 is much more severe, we postulate that loss of only RPN-10, RPN-13, or DSS-1 may not sufficiently impair the ability of the other subunits to feed substrates to RPN-1 for movement through the base and into the proteasome core. Takahashi *et al.* previously showed redundancy between *rpn-10* and *rpn-12* (Takahashi *et al.*, 2002). Structural analyses place RPN-10 at the interface of the 19S base and lid, linking RPN-1 to RPN-12 (see Figure 1A). In the absence of RPN-10, these two subunits may directly interact, as suggested by dynamic models of proteasome structure with and without substrate (Bard *et al.*, 2018). Alternatively, however, these data may suggest that the 19S lid adopts a novel structure in the worm germ line. Existence of tissue-specific proteasomes is not unprecedented but the study of these variants is still into its infancy (Kish-Trier and Hill, 2013; Uechi, Hamazaki and Murata, 2014; Gómez-H *et al.*, 2019; Motosugi and Murata, 2019). These modified proteasomes provide a mechanism to adapt to tissue-specific needs. Determining whether the *C. elegans* 19S RP adopts a germ line specific configuration is an important avenue for future investigation.

Most 19S RP subunit depletions caused aberrant nuclear accumulation of GFP::WEE-1.3. However, bortezomib treatment did not alter the localization of WEE-1.3. Bortezomib works by binding to the β5 subunit of the 20S CP and inhibiting its peptidase activity, whereas depletion of specific 19S subunits may weaken 19S RP and 20S CP interactions, destabilizing part or all of the proteasome structure or may impair 19S RP-substrate interactions (Adams *et al.*, 1999; Bai *et al.*, 2019; Thibaudeau and Smith, 2019). Therefore, we speculate that an intact, stable proteasome structure, but not its activity, is required for the proper perinuclear localization of WEE-1.3 (Figure 8B). While proteolytic roles of the proteasome are well established, growing evidence supports additional roles for intact proteasome (or its subcomplexes), including in the cell cycle, transcription, and chromatin organization (Nishiyama *et al.*, 2000; Geng, Wenzel and Tansey, 2012; Seo *et al.*, 2017). One possibility is that the proteasome tethers WEE-1.3 to the perinuclear region, potentially even the nuclear pore complex, through protein-protein interactions (Albert *et al.*, 2017).

Our studies also point to differences between the behavior of the 19S lid and base. With exception of *rpt-2*, none of 19S base subunits were able to suppress *wee-1(RNAi)*-induced sterility, whereas many of the lid subunits did suppress. These data could be explained if the lid has independent, non-proteasomal functions or that it combines with other proteins to make an alternative regulatory particle. In favor of the former model, we previously showed that proteasome inhibition by bortezomib failed to suppress *wee-1.3(RNAi)* infertility suggesting that the misregulation of protein turnover is not driving the oocyte maturation defect of *wee-1.3* depletion (Fernando, Elliot and Allen, 2020). The mechanism by which the suppression of *wee-1.3(RNAi)* infertility occurs is still unknown but future studies may offer new insights into the regulation of this highly conserved WEE-1.3/Myt1 cell cycle kinase.

Previous research in *C. elegans* showed that RPT-6 has a role in transcription. RPT-6 interacts with the transcription factor ELT-2 to regulate expression of immune response genes and this role is independent of the proteolytic activity of the proteasome (Olaitan and Aballay, 2018). Therefore, our observation that depletion of RPT-6 does not affect germline proteolytic function, but rather causes a reduced brood and larval arrest can mean two things: either RPT-6 is a developmental stage specific proteasome subunit that is essential for proteolytic function of the proteasome only during larval development; or, RPT-6 may play non-proteolytic roles in the *C. elegans* germ line because depletion of RPT-6 causes a reduced brood but overall germ line proteolytic function is not affected. While we favor, off-proteasome functions for RPT-6 in controlling oocyte quality, further studies are needed to elucidate RPT-6 function. It is noteworthy that RPT-6 is known to play non-proteolytic roles in transcription in both yeast and mammalian cells (Chang *et al.*, 2001; Gonzalez *et al.*, 2002; Lee *et al.*, 2005; Uprety *et al.*, 2012).

Endogenous GFP tagging of a number of the 19S proteasomal subunits indicated strong expression throughout the germ line of *C. elegans*, in addition to ubiquitous, somatic expression. However, we are the first to report isoform-specific localization of RPN-6.1 in *C. elegans*. With isoform RPN-6.1A being expressed only in the body wall muscles while RPN-6.1::OLLAS (which marks both Isoforms A and B) is expressed throughout the hermaphrodite female germ line but is distinctly absent from both sperm and the pharynx. Since downregulation of RPN-6.1 causes severe dysfunction of the proteolytic activity of the proteasome, we speculate that there is likely to be other RPN-6.1 variant(s) that functions in the pharynx and sperm (Vilchez, Morantte, *et al.*, 2012; Fernando, Elliot and Allen, 2020). Indeed, RPN-6.2, a RPN-6 paralog, has recently been identified as sperm-specific (personal communication, Lynn Boyd). Sperm-specific proteasome subunits have been described in various systems and may exist to meet the massive protein turnover for the histone to protamine transition or to facilitate fertilization (Belote and Zhong, 2009; Sutovsky, 2011; Uechi, Hamazaki and Murata, 2014; Zhang *et al.*, 2019; Palacios *et al.*, 2021). One critical remaining question is whether the different isoforms reflect tissue-specific modifications or adaptations to specific substrate in these tissues. Further analysis of these questions in the worm will enhance our knowledge of the diverse and dynamic regulation of the proteasome in different tissues.

RPN-6.1/Rpn6/PSMD11 is one of the subunits known to play a crucial role in proteasome stability and lid subcomplex assembly (Santamaría *et al.*, 2003; Isono *et al.*, 2005; Bai *et al.*, 2019). Our results suggest that *C. elegans* RPN-6.1 and RPN-7 aid in the nuclear localization of the lid subcomplex. Our future studies will focus on determining the mechanism by which RPN-6.1 and RPN-7 aid in this process. Interestingly, neither RPN-6.1 nor RPN-7 possess canonical NLS sequences, implying either the proteins have cryptic NLSs or that additional binding partners are required for nuclear localization of the 19S RP lid subcomplexes. The endogenously-tagged strains that we generated will be beneficial in both biochemical and genetic experiments to identify such sequences or chaperones binding partners. Obtaining a complete set of fluorescently tagged lid subunits will aid in further elucidating the mechanism by which the lid subcomplex assembles and becomes nuclear localized using the *C. elegans* germ line as a model system.

The spatiotemporal and depletion analyses of the *C. elegans* proteasome subunits in this study reveal differential roles being played by specific subunits and provides crucial information to fill the knowledge gaps in our understanding of the 26S proteasome and its many functions. Generation of these endogenously fluorescently tagged 19S RP subunits and future tagged subunits will serve as valuable resources for future proteasome subunits. Our current findings in the multicellular model *C. elegans* and the future ones that stem from this research have tremendous potential to transform the proteasome field and can be translated into better understanding human proteasome function.

## Author Contributions

LMF and CQC designed experiments, performed experiments and wrote aspects of the manuscript. MM and CU performed experiments. AKA and JLY conceived the projects, designed experiments, and wrote aspects of the manuscript. All authors revised the manuscript.

## Acknowledgements

We thank the undergraduate researchers in the Allen lab for their assistance with general laboratory tasks that enabled this research; the Duttaroy and Robinson Labs at Howard University for sharing equipment and reagents; Dr. Kuppuswamy Subramaniam for kindly providing the IT1187 strain; and members of the Baltimore Worm Club for helpful discussions. We also thank Dr. Valentin Boerner and his laboratory for helpful discussions and all members of the Yanowitz for input into the experiments.

## References

Adams, J. et al. (1999) ‘Proteasome inhibitors: A novel class of potent and effective antitumor agents’, Cancer Research, 59(11).

Ahuja, J. S. et al. (2017) ‘Control of meiotic pairing and recombination by chromosomally tethered 26S proteasome’, Science, 355(6323). doi: 10.1126/science.aaf4778.

Albert, S. et al. (2017) ‘Proteasomes tether to two distinct sites at the nuclear pore complex’, Proceedings of the National Academy of Sciences of the United States of America, 114(52). doi: 10.1073/pnas.1716305114.

Allen, A. K., Nesmith, J. E. and Golden, A. (2014) ‘An RNAi-based suppressor screen identifies interactors of the Myt1 ortholog of *Caenorhabditis elegans*.’, G3 (Bethesda, Md.). United States, 4(12), pp. 2329–2343. doi: 10.1534/g3.114.013649.

Arribere, J. A. et al. (2014) ‘Efficient marker-free recovery of custom genetic modifications with CRISPR/Cas9 in *Caenorhabditis elegans*’, Genetics, 198(3), pp. 837–846. doi: 10.1534/genetics.114.169730.

Bai, M. et al. (2019) ‘In-depth analysis of the lid subunits assembly mechanism in mammals’, Biomolecules. doi: 10.3390/biom9060213.

Bard, J. A. M. et al. (2018) ‘Structure and Function of the 26S Proteasome’, Annual Review of Biochemistry. doi: 10.1146/annurev-biochem-062917-011931.

Beck, F. et al. (2012) ‘Near-atomic resolution structural model of the yeast 26S proteasome.’, Proceedings of the National Academy of Sciences of the United States of America, 109(37), pp. 14870–14875. doi: 10.1073/pnas.1213333109.

Belote, J. M. and Zhong, L. (2009) ‘Duplicated proteasome subunit genes in Drosophila and their roles in spermatogenesis’, Heredity, 103(1). doi: 10.1038/hdy.2009.23.

Bhat, K. P. and Greer, S. F. (2011) ‘Proteolytic and non-proteolytic roles of ubiquitin and the ubiquitin proteasome system in transcriptional regulation’, Biochimica et Biophysica Acta - Gene Regulatory Mechanisms, pp. 150–155. doi: 10.1016/j.bbagrm.2010.11.006.

Boateng, R. et al. (2017) ‘Novel functions for the RNA-binding protein ETR-1 in *Caenorhabditis* elegans reproduction and engulfment of germline apoptotic cell corpses’, Developmental Biology. doi: 10.1016/j.ydbio.2017.06.015.

Brenner, S. (1974) ‘The genetics of *Caenorhabditis elegans*’, Genetics. 1974/05/01, 77(1), pp. 71–94. Available at: http://www.ncbi.nlm.nih.gov/entrez/query.fcgi?cmd=Retrieve&db=PubMed&dopt=Citation&list_uids=4366476.

Brooks, P. et al. (2000) ‘Subcellular localization of proteasomes and their regulatory complexes in mammalian cells.’, The Biochemical journal, 346 Pt 1, pp. 155–161. doi: 10.1042/0264-6021:3460155.

Budenholzer, L. et al. (2017) ‘Proteasome Structure and Assembly’, Journal of Molecular Biology, pp. 3500–3524. doi: 10.1016/j.jmb.2017.05.027.

Budenholzer, L. et al. (2020) ‘The Sts1 nuclear import adapter uses a non-canonical bipartite nuclear localization signal and is directly degraded by the proteasome’, Journal of Cell Science, 133(6), p. jcs236158. doi: 10.1242/jcs.236158.

Burrows, A. E. et al. (2006) ‘The *C. elegans* Myt1 ortholog is required for the proper timing of oocyte maturation’, Development. 2006/01/20, 133(4), pp. 697–709. doi: dev.02241 [pii] 10.1242/dev.02241.

Chang, C. et al. (2001) ‘The Gal4 Activation Domain Binds Sug2 Protein, a Proteasome Component, in Vivo and in Vitro’, Journal of Biological Chemistry, 276(33). doi: 10.1074/jbc.M102254200.

Chen, J. J. and Arur, S. (2017) ‘Discovering Functional ERK Substrates Regulating *Caenorhabditis elegans* Germline Development’, Methods in molecular biology (Clifton, N.J.), 1487, pp. 317–335. doi: 10.1007/978-1-4939-6424-6_24.

Colaiácovo, M. P. et al. (2003) ‘Synaptonemal complex assembly in *C. elegans* is dispensable for loading strand-exchange proteins but critical for proper completion of recombination’, Developmental Cell, 5(3). doi: 10.1016/S1534-5807(03)00232-6.

Couteau, F. and Zetka, M. (2005) ‘HTP-1 coordinates synaptonemal complex assembly with homolog alignment during meiosis in *C. elegans*’, Genes and Development, 19(22). doi: 10.1101/gad.1348205.

Crittenden, S. L. et al. (2006) ‘Cellular analyses of the mitotic region in the *Caenorhabditis elegans* adult germ line.’, Molecular biology of the cell, 17(7), pp. 3051–61. doi: 10.1091/mbc.E06-03-0170.

Dantuma, N. P. et al. (2000) ‘Short-lived green fluorescent proteins for quantifying ubiquitin/proteasome-dependent proteolysis in living cells.’, Nature biotechnology. United States, 18(5), pp. 538–543. doi: 10.1038/75406.

Enenkel, C. (2014) ‘Nuclear transport of yeast proteasomes’, Biomolecules. MDPI, 4(4), pp. 940–955. doi: 10.3390/biom4040940.

Ferdous, A., Kodadek, T. and Johnston, S. A. (2002) ‘A nonproteolytic function of the 19S regulatory subunit of the 26S proteasome is required for efficient activated transcription by human RNA polymerase II.’, Biochemistry, 41(42), pp. 12798–805. doi: 10.1021/bi020425t.

Fernando, L. M., Elliot, J. and Allen, A. K. (2020) ‘The *Caenorhabditis elegans* proteasome subunit RPN **‐**12 is required for hermaphrodite germline sex determination and oocyte quality.’, Developmental Dynamics. doi: 10.1002/dvdy.235.

Finley, D. (2009) ‘Recognition and Processing of Ubiquitin-Protein Conjugates by the Proteasome’, Annu Rev Biochem, 78, pp. 477–513. doi: 10.1146/annurev.biochem.78.081507.101607.Recognition.

Funakoshi, M. et al. (2009) ‘Multiple assembly chaperones govern biogenesis of the proteasome regulatory particle base’, Cell. 2009/05/14, 137(5), pp. 887–899. doi: 10.1016/j.cell.2009.04.061.

Geng, F., Wenzel, S. and Tansey, W. P. (2012) ‘Ubiquitin and proteasomes in transcription’, Annual Review of Biochemistry, 81. doi: 10.1146/annurev-biochem-052110-120012.

Glotzer, M., Murray, A. W. and Kirschner, M. W. (1991) ‘Cyclin is degraded by the ubiquitin pathway.’, Nature, 349(6305), pp. 132–8. doi: 10.1038/349132a0.

Goldstein, P. (1986) ‘The synaptonemal complexes of *Caenorhabditis elegans*: the dominant him mutant mnT6 and pachytene karyotype analysis of the X-autosome translocation’, Chromosoma, 93(3). doi: 10.1007/BF00292746.

Gómez-H, L. et al. (2019) ‘The PSMA8 subunit of the spermatoproteasome is essential for proper meiotic exit and mouse fertility.’, PLoS genetics, 15(8), p. e1008316. doi: 10.1371/journal.pgen.1008316.

Gonzalez, F. et al. (2002) ‘Recruitment of a 19S proteasome subcomplex to an activated promoter.’, Science (New York, N.Y.), 296(5567), pp. 548–50. doi: 10.1126/science.1069490.

Greenstein, D. (2005) ‘Control of oocyte meiotic maturation and fertilization.’, WormBook : the online review of C. elegans biology, pp. 1–12. doi: 10.1895/wormbook.1.53.1.

Groll, M. et al. (1997) ‘Structure of 20S proteasome from yeast at 2.4Å resolution’, Nature, 386(6624), pp. 463–471. doi: 10.1038/386463a0.

Hamazaki, J., Hirayama, S. and Murata, S. (2015) ‘Redundant Roles of Rpn10 and Rpn13 in Recognition of Ubiquitinated Proteins and Cellular Homeostasis’, PLoS Genetics, 11(7). doi: 10.1371/journal.pgen.1005401.

Hamer, G., Matilainen, O. and Holmberg, C. I. (2010) ‘A photoconvertible reporter of the ubiquitin-proteasome system in vivo’, Nature methods, 7(6), pp. 473–8. doi: 10.1038/nmeth.1460.

Hanna, J. and Finley, D. (2007) ‘A proteasome for all occasions’, FEBS Letters, pp. 2854–2861. doi: 10.1016/j.febslet.2007.03.053.

Hillers, K. J. et al. (2015) ‘Meiosis.’, WormBook : the online review of C. elegans biology, pp. 1–54. doi: 10.1895/wormbook.1.178.1.

Hirano, Y. et al. (2006) ‘Cooperation of Multiple Chaperones Required for the Assembly of Mammalian 20S Proteasomes’, Molecular Cell, 24(6). doi: 10.1016/j.molcel.2006.11.015.

Hirano, Y. et al. (2008) ‘Dissecting beta-ring assembly pathway of the mammalian 20S proteasome’, The EMBO journal. 2008/07/24. Nature Publishing Group, 27(16), pp. 2204–2213. doi: 10.1038/emboj.2008.148.

Hochstrasser, M. (1996) ‘Ubiquitin-dependent protein degradation’, Annual review of genetics, 30(1), pp. 405–439. doi: 10.1146/annurev.genet.30.1.405.

Huang, X. et al. (2016) ‘An atomic structure of the human 26S proteasome’, Nature Structural & Molecular Biology, 23(9), pp. 778–785. doi: 10.1038/nsmb.3273.

Hubbard, E. J. A. (2007) ‘*Caenorhabditis elegans* germ line: a model for stem cell biology.’, Developmental dynamics, 236(12), pp. 3343–57. doi: 10.1002/dvdy.21335.

Hubbard, E. J. and Greenstein, D. (2000) ‘The *Caenorhabditis elegans* gonad: a test tube for cell and developmental biology’, Dev Dyn, 218(1), pp. 2–22. Available at: http://www.ncbi.nlm.nih.gov/entrez/query.fcgi?cmd=Retrieve&db=PubMed&dopt=Citation&list_uids=10822256.

Inoue, T. et al. (2006) ‘Cell cycle control by daf-21/Hsp90 at the first meiotic prophase/metaphase boundary during oogenesis in *Caenorhabditis elegans*’, Dev Growth Differ. 2006/02/10, 48(1), pp. 25–32. doi: 10.1111/j.1440-169X.2006.00841.x.

Isono, E. et al. (2005) ‘Functional analysis of Rpn6p, a lid component of the 26 S proteasome, using temperature-sensitive rpn6 mutants of the yeast *Saccharomyces cerevisiae*’, Journal of Biological Chemistry, 280(8). doi: 10.1074/jbc.M409364200.

Isono, E. et al. (2007) ‘The assembly pathway of the 19S regulatory particle of the yeast 26S proteasome’, Molecular biology of the cell. 2006/11/29. The American Society for Cell Biology, 18(2), pp. 569–580. doi: 10.1091/mbc.e06-07-0635.

Kaneko, T. et al. (2009) ‘Assembly pathway of the Mammalian proteasome base subcomplex is mediated by multiple specific chaperones.’, Cell. United States, 137(5), pp. 914–925. doi: 10.1016/j.cell.2009.05.008.

Kim, H. M., Yu, Y. and Cheng, Y. (2011) ‘Structure characterization of the 26S proteasome.’, Biochimica et biophysica acta, 1809(2), pp. 67–79. doi: 10.1016/j.bbagrm.2010.08.008.

Kish-Trier, E. and Hill, C. P. (2013) ‘Structural Biology of the Proteasome’, Annual Review of Biophysics, 42, pp. 29–49. doi: 10.1146/annurev-biophys-083012-130417.

Kocsisova, Z., Kornfeld, K. and Schedl, T. (2019) ‘Rapid population-wide declines in stem cell number and activity during reproductive aging in *C. elegans*’, Development (Cambridge), 146(8). doi: 10.1242/dev.173195.

Kumar, G. A. and Subramaniam, K. (2018) ‘PUF-8 facilitates homologous chromosome pairing by promoting proteasome activity during meiotic entry *in C. elegans*’, Development (Cambridge). England, 145(7). doi: 10.1242/dev.163949.

Kusmierczyk, A. R. et al. (2008) ‘A multimeric assembly factor controls the formation of alternative 20S proteasomes’, Nature Structural and Molecular Biology, 15(3). doi: 10.1038/nsmb.1389.

Kwak, J., Workman, J. L. and Lee, D. (2011) ‘The proteasome and its regulatory roles in gene expression’, Biochimica et Biophysica Acta - Gene Regulatory Mechanisms, pp. 88–96. doi: 10.1016/j.bbagrm.2010.08.001.

Lamitina, S. T. and L’Hernault, S. W. (2002) ‘Dominant mutations in the *Caenorhabditis elegans* Myt1 ortholog *wee-1.3* reveal a novel domain that controls M-phase entry during spermatogenesis’, Development, 129(21), pp. 5009–5018. Available at: http://www.ncbi.nlm.nih.gov/entrez/query.fcgi?cmd=Retrieve&db=PubMed&dopt=Citation&list_uids=12397109.

Lee, D. et al. (2005) ‘The proteasome regulatory particle alters the SAGA coactivator to enhance its interactions with transcriptional activators’, Cell, 123(3). doi: 10.1016/j.cell.2005.08.015.

Lee, M. H. and Schedl, T. (2010) ‘*C. elegans* STAR proteins, GLD-1 and ASD-2, regulate specific RNA targets to control development’, Advances in Experimental Medicine and Biology, 693, pp. 106–122. doi: 10.1007/978-1-4419-7005-3_8.

Lehmann, A. et al. (2002) ‘20 S proteasomes are imported as precursor complexes into the nucleus of yeast.’ Edited by R. Huber’, Journal of Molecular Biology, 317(3), pp. 401–413. doi: https://doi.org/10.1006/jmbi.2002.5443.

Li, X. et al. (2007) ‘beta-Subunit appendages promote 20S proteasome assembly by overcoming an Ump1-dependent checkpoint’, The EMBO journal. 2007/04/12. Nature Publishing Group, 26(9), pp. 2339–2349. doi: 10.1038/sj.emboj.7601681.

Li, X. et al. (2013) ‘Electron counting and beam-induced motion correction enable near-atomic-resolution single-particle cryo-EM’, Nature Methods, 10(6), pp. 584–590. doi: 10.1038/nmeth.2472.

Li, Y., Li, S. and Wu, H. (2022) ‘Ubiquitination-Proteasome System (UPS) and Autophagy Two Main Protein Degradation Machineries in Response to Cell Stress’, Cells, 11(5). doi: 10.3390/cells11050851.

Liu, H. et al. (2012) ‘Enhancement of 26S proteasome functionality connects oxidative stress and vascular endothelial inflammatory response in diabetes mellitus’, Arteriosclerosis, Thrombosis, and Vascular Biology. doi: 10.1161/ATVBAHA.112.253385.

Lokireddy, S., Kukushkin, N. V. and Goldberg, A. L. (2015) ‘cAMP-induced phosphorylation of 26S proteasomes on Rpn6/PSMD11 enhances their activity and the degradation of misfolded proteins.’, Proceedings of the National Academy of Sciences of the United States of America, 112(52), pp. E7176–85. doi: 10.1073/pnas.1522332112.

MacQueen, A. J. et al. (2002) ‘Synapsis-dependent and -independent mechanisms stabilize homolog pairing during meiotic prophase in *C. elegans*’, Genes and Development, 16(18). doi: 10.1101/gad.1011602.

Mainpal, R., Nance, J. and Yanowitz, J. L. (2015) ‘A germ cell determinant reveals parallel pathways for germ line development in *Caenorhabditis elegans*’, Development (Cambridge), 142(20). doi: 10.1242/dev.125732.

Maneix, L. and Catic, A. (2016) ‘Touch and go: Nuclear proteolysis in the regulation of metabolic genes and cancer’, FEBS Letters, pp. 908–923. doi: 10.1002/1873-3468.12087.

Marshall, R. S. and Vierstra, R. D. (2019) ‘Dynamic regulation of the 26S proteasome: From synthesis to degradation’, Frontiers in Molecular Biosciences. doi: 10.3389/fmolb.2019.00040.

Martinez-Perez, E. and Villeneuve, A. M. (2005) ‘HTP-1-dependent constraints coordinate homolog pairing and synapsis and promote chiasma formation during *C. elegans* meiosis’, Genes and Development, 19(22). doi: 10.1101/gad.1338505.

Mikkonen, E., Haglund, C. and Holmberg, C. I. (2017) ‘Immunohistochemical analysis reveals variations in proteasome tissue expression in *C. elegans*’, PLoS ONE, 12(8). doi: 10.1371/journal.pone.0183403.

Motosugi, R. and Murata, S. (2019) ‘Dynamic regulation of proteasome expression’, Frontiers in Molecular Biosciences, 6(MAY), pp. 4–11. doi: 10.3389/fmolb.2019.00030.

Murata, S., Yashiroda, H. and Tanaka, K. (2009) ‘Molecular mechanisms of proteasome assembly’, Nature Reviews Molecular Cell Biology. doi: 10.1038/nrm2630.

Myeku, N. et al. (2011) ‘Assessment of proteasome impairment and accumulation/aggregation of ubiquitinated proteins in neuronal cultures’, Methods in Molecular Biology. doi: 10.1007/978-1-61779-328-8_18.

Nishiyama, A. et al. (2000) ‘A nonproteolytic function of the proteasome is required for the dissociation of CDc2 and cyclin B at the end of M phase’, Genes and Development, 14(18), pp. 2344–2357. doi: 10.1101/gad.823200.

Olaitan, A. O. and Aballay, A. (2018) ‘Non-proteolytic activity of 19S proteasome subunit RPT-6 regulates GATA transcription during response to infection’, PLoS Genetics, 14(9). doi: 10.1371/journal.pgen.1007693.

Pack, C. G. et al. (2014) ‘Quantitative live-cell imaging reveals spatio-temporal dynamics and cytoplasmic assembly of the 26S proteasome’, Nature Communications, 5. doi: 10.1038/ncomms4396.

Paix, A. et al. (2015) ‘High Efficiency, Homology-Directed Genome Editing in Caenorhabditis elegans Using CRISPR/Cas9 Ribonucleoprotein Complexes.’, Genetics. doi: 10.1534/genetics.115.179382.

Palacios, V. et al. (2021) ‘Importin-9 regulates chromosome segregation and packaging in Drosophila germ cells’, Journal of Cell Science, 134(7). doi: 10.1242/jcs.258391.

Paraskevopoulos, K. et al. (2014) ‘Dss1 is a 26S proteasome ubiquitin receptor’, Molecular cell. 2014/10/09. Cell Press, 56(3), pp. 453–461. doi: 10.1016/j.molcel.2014.09.008.

Pispa, J. et al. (2008) ‘*C. elegans dss-1* is functionally conserved and required for oogenesis and larval growth’, BMC Dev Biol, 8, p. 51. doi: 1471-213X-8-51 [pii]\r 10.1186/1471-213X-8-51.

Pispa, J., Matilainen, O. and Holmberg, C. I. (2020) ‘Tissue-specific effects of temperature on proteasome function’, Cell stress & chaperones. 2020/04/18. Springer Netherlands, 25(3), pp. 563–572. doi: 10.1007/s12192-020-01107-y.

Prasada Rao, H. B. D. et al. (2017) ‘A SUMO-ubiquitin relay recruits proteasomes to chromosome axes to regulate meiotic recombination’, Science, 355(6323). doi: 10.1126/science.aaf6407.

Roelofs, J. et al. (2009) ‘Chaperone-mediated pathway of proteasome regulatory particle assembly’, Nature, 459(7248), pp. 861–865. doi: 10.1038/nature08063.

Rog, O., Köhler, S. and Dernburg, A. F. (2017) ‘The synaptonemal complex has liquid crystalline properties and spatially regulates meiotic recombination factors’, eLife, 6. doi: 10.7554/eLife.21455.

Ruiz, E. J., Vilar, M. and Nebreda, A. R. (2010) ‘A two-step inactivation mechanism of Myt1 ensures CDK1/cyclin B activation and meiosis I entry.’, Current Biology. 2010/04/07, 20(8), pp. 717–23. doi: 10.1016/j.cub.2010.02.050.

Saez, I. and Vilchez, D. (2014) ‘The mechanistic links between proteasome activity, aging and age-related diseases.’, Current genomics, 15(1), pp. 38–51. doi: 10.2174/138920291501140306113344.

Santamaría, P. G. et al. (2003) ‘Rpn6p, a proteasome subunit from *Saccharomyces cerevisiae*, is essential for the assembly and activity of the 26 S proteasome’, Journal of Biological Chemistry, 278(9). doi: 10.1074/jbc.M209420200.

Satoh, K. et al. (2001) ‘Assembly of the 26S proteasome is regulated by phosphorylation of the p45/Rpt6 ATPase subunit’, Biochemistry, 40(2), pp. 314–319. doi: 10.1021/bi001815n.

Schmidt, M. and Finley, D. (2014) ‘Regulation of proteasome activity in health and disease.’, Biochimica biophysica acta, 1843(1), pp. 13–25. doi: 10.1016/j.bbamcr.2013.08.012.

Seo, H. D. et al. (2017) ‘The 19S proteasome is directly involved in the regulation of heterochromatin spreading in fission yeast’, Journal of Biological Chemistry, 292(41), pp. 17144–17155. doi: 10.1074/jbc.M117.790824.

Shimada, M. et al. (2006) ‘Proteasomal ubiquitin receptor RPN-10 controls sex determination in *Caenorhabditis elegans*’, Molecular Biology of the Cell, 17(12), pp. 5356–5371. doi: 10.1091/mbc.E06-05-0437.

Silva, N. et al. (2014) ‘The fidelity of synaptonemal complex assembly is regulated by a signaling mechanism that controls early meiotic progression’, Developmental Cell, 31(4). doi: 10.1016/j.devcel.2014.10.001.

Sutovsky, P. (2011) ‘Sperm proteasome and fertilization’, Reproduction. doi: 10.1530/REP-11-0041.

Takahashi, M. et al. (2002) ‘Reverse genetic analysis of the *Caenorhabditis elegans* 26S proteasome subunits by RNA interference’, Biological Chemistry, 383(7–8), pp. 1263–1266. doi: 10.1515/BC.2002.140.

Le Tallec, B. et al. (2007) ‘20S proteasome assembly is orchestrated by two distinct pairs of chaperones in yeast and in mammals.’, Molecular cell. United States, 27(4), pp. 660–674. doi: 10.1016/j.molcel.2007.06.025.

Tanaka, K. et al. (1990) ‘Possible mechanism of nuclear translocation of proteasomes’, FEBS Letters. John Wiley & Sons, Ltd, 271(1–2), pp. 41–46. doi: https://doi.org/10.1016/0014-5793(90)80367-R.

Thibaudeau, T. A. and Smith, D. M. (2019) ‘A practical review of proteasome pharmacology’, Pharmacological Reviews, 71(2). doi: 10.1124/pr.117.015370.

Timmons, L., Court, D. L. and Fire, a (2001) ‘Ingestion of bacterially expressed dsRNAs can produce specific and potent genetic interference in *Caenorhabditis elegans*.’, Gene, 263(1–2), pp. 103–12. Available at: http://www.ncbi.nlm.nih.gov/pubmed/11223248.

Tomko, R. J. and Hochstrasser, M. (2011) ‘Incorporation of the Rpn12 Subunit Couples Completion of Proteasome Regulatory Particle Lid Assembly to Lid-Base Joining’, Molecular Cell. doi: 10.1016/j.molcel.2011.11.020.

Uechi, H., Hamazaki, J. and Murata, S. (2014) ‘Characterization of the testis-specific proteasome subunit α4s in mammals’, The Journal of biological chemistry. 2014/03/25. American Society for Biochemistry and Molecular Biology, 289(18), pp. 12365–12374. doi: 10.1074/jbc.M114.558866.

Unno, M. et al. (2002) ‘The Structure of the Mammalian 20S Proteasome at 2.75 Å Resolution’, Structure, 10(5), pp. 609–618. doi: https://doi.org/10.1016/S0969-2126(02)00748-7.

Uprety, B. et al. (2012) ‘The 19S proteasome subcomplex promotes the targeting of NuA4 HAT to the promoters of ribosomal protein genes to facilitate the recruitment of TFIID for transcriptional initiation in vivo’, Nucleic Acids Research. doi: 10.1093/nar/gkr977.

Vilchez, D., Boyer, L., et al. (2012) ‘Increased proteasome activity in human embryonic stem cells is regulated by PSMD11’, Nature. doi: 10.1038/nature11468.

Vilchez, D., Morantte, I., et al. (2012) ‘RPN-6 determines *C. elegans* longevity under proteotoxic stress conditions.’, Nature. Nature Publishing Group, 489(7415), pp. 263–8. doi: 10.1038/nature11315.

Wagner, C. R. et al. (2010) ‘Xnd-1 regulates the global recombination landscape in *Caenorhabditis elegans*’, Nature, 467(7317). doi: 10.1038/nature09429.

Walerych, D. et al. (2016) ‘Proteasome machinery is instrumental in a common gain-of-function program of the p53 missense mutants in cancer’, Nature Cell Biology, 18, pp. 897–909. doi: 10.1038/ncb3380.

Wendler, P. et al. (2004) ‘The bipartite nuclear localization sequence of Rpn2 is required for nuclear import of proteasomal base complexes via karyopherin alphabeta and proteasome functions.’, The Journal of biological chemistry. United States, 279(36), pp. 37751–37762. doi: 10.1074/jbc.M403551200.

Wendler, P. and Enenkel, C. (2019) ‘Nuclear transport of yeast proteasomes’, Frontiers in Molecular Biosciences, 6(MAY), pp. 1–4. doi: 10.3389/fmolb.2019.00034.

Wormbase. (2022) http://www.wormbase.org/db/get?name=WBGene00004462;class=gene

Wu, W. et al. (2018) ‘PAC1-PAC2 proteasome assembly chaperone retains the core α4–α7 assembly intermediates in the cytoplasm’, Genes to Cells, 23(10). doi: 10.1111/gtc.12631.

Yashiroda, H. et al. (2008) ‘Crystal structure of a chaperone complex that contributes to the assembly of yeast 20S proteasomes.’, Nature structural & molecular biology. United States, 15(3), pp. 228–236. doi: 10.1038/nsmb.1386.

Zhang, Q. et al. (2019) ‘Meiosis I progression in spermatogenesis requires a type of testis-specific 20S core proteasome’, Nature Communications, 10(1). doi: 10.1038/s41467-019-11346-y.

Zou, T. and Lin, Z. (2021) ‘The involvement of ubiquitination machinery in cell cycle regulation and cancer progression’, International Journal of Molecular Sciences. doi: 10.3390/ijms22115754.

