## Supplemental Files for "Proteasomal subunit depletions differentially affect germline integrity in *C. elegans*"

**Supplemental Figure 1**

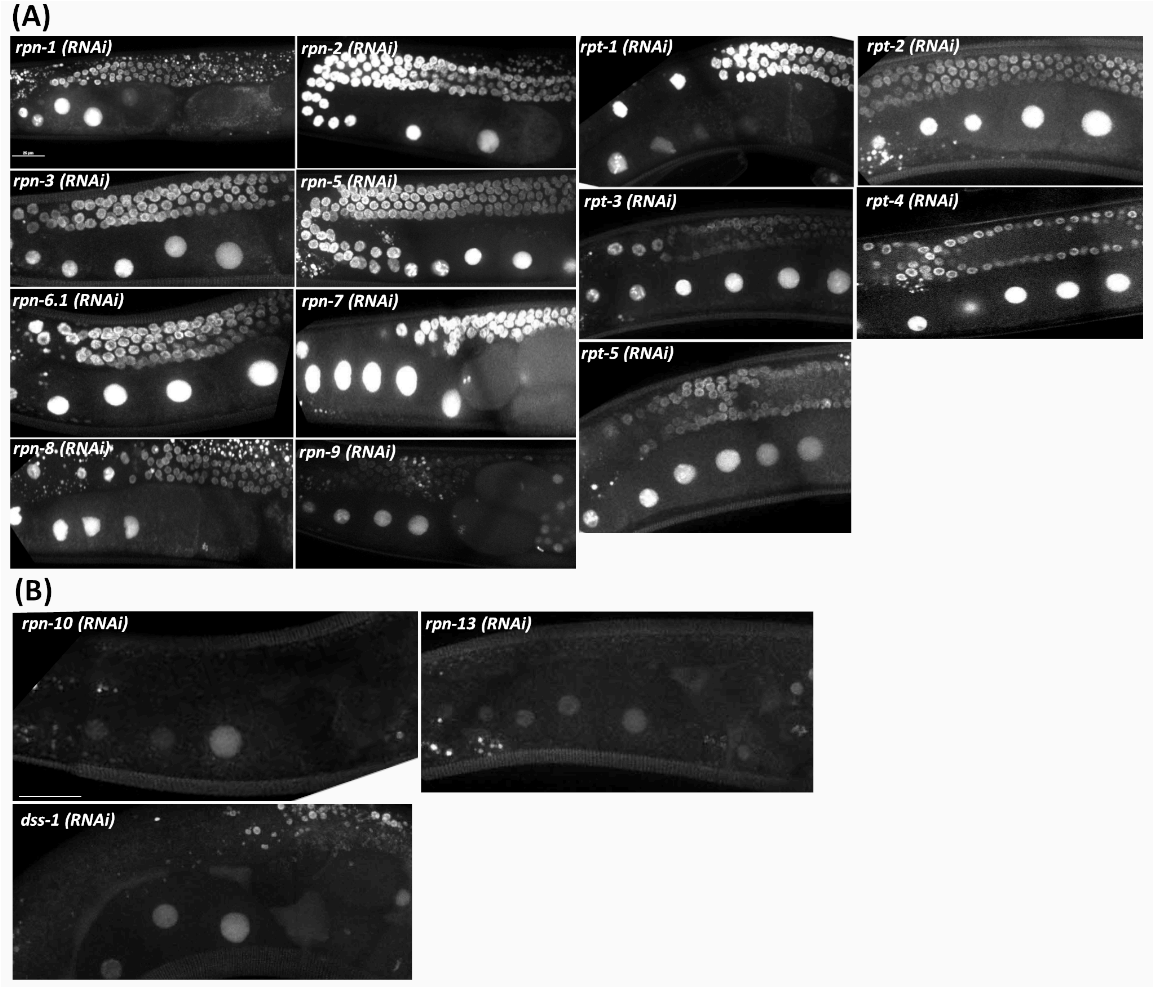


**Supplemental Figure 1. Ub(G76V)::GFP::H2B animals RNA-depleted of 19S RP subunits to assess the germline proteolytic activity of the proteasome**. (A) Live imaging of germ lines from Ub(G76V)::GFP::H2B animals RNAi-depleted of the indicated 19S RP subunits of the 26S proteasome. The increase in the fluorescence in Ub(G76V)::GFP::H2B germ lines indicates reduced proteolytic activity of the proteasome. (B) RNAi depletion of RPN-10, RPN-13 and DSS-1 did not increase fluorescence in Ub(G76V)::GFP::H2B germ line suggesting proper function of the proteasome.

**Supplemental Figure 2**

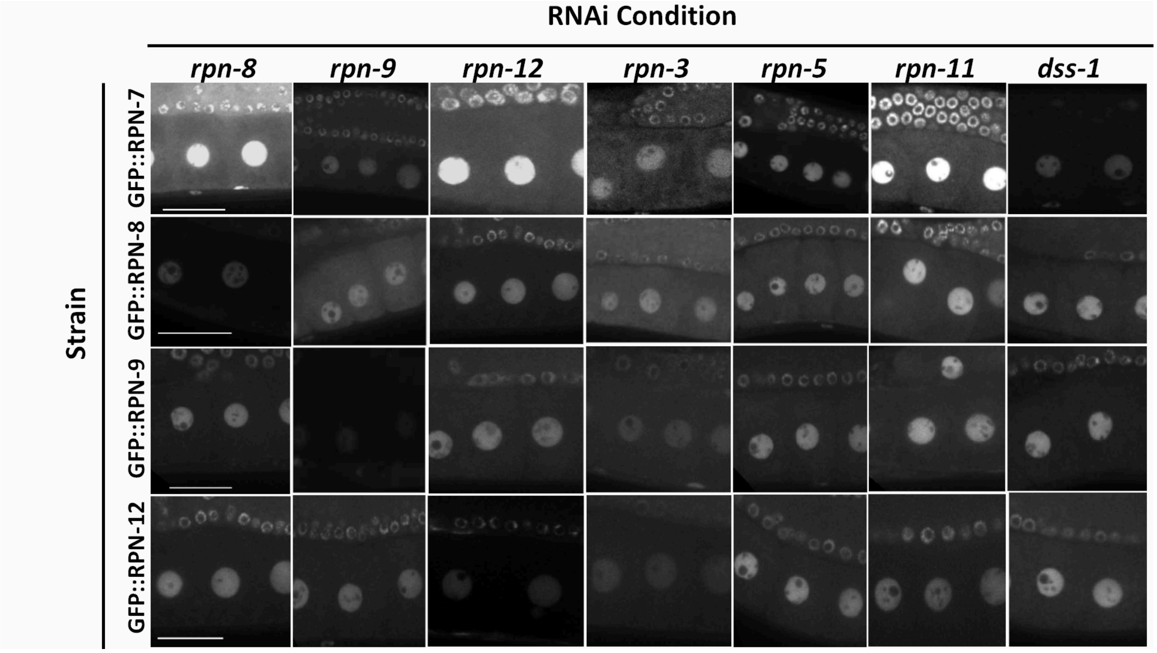


**Supplemental Figure 2. Localization of specific 19S RP lid subunits under various lid subunit RNAi conditions.** Depletion of RPN-8, RPN-9, RPN-12, RPN-3, RPN-5, RPN-11 and DSS-1 via RNAi in GFP::RPN-7, GFP::RPN-8, GFP::RPN-9 and GFP::RPN-12 expressing oocytes (n=15-20). Scale bar represents 25μm.

**Supplemental Figure 3**

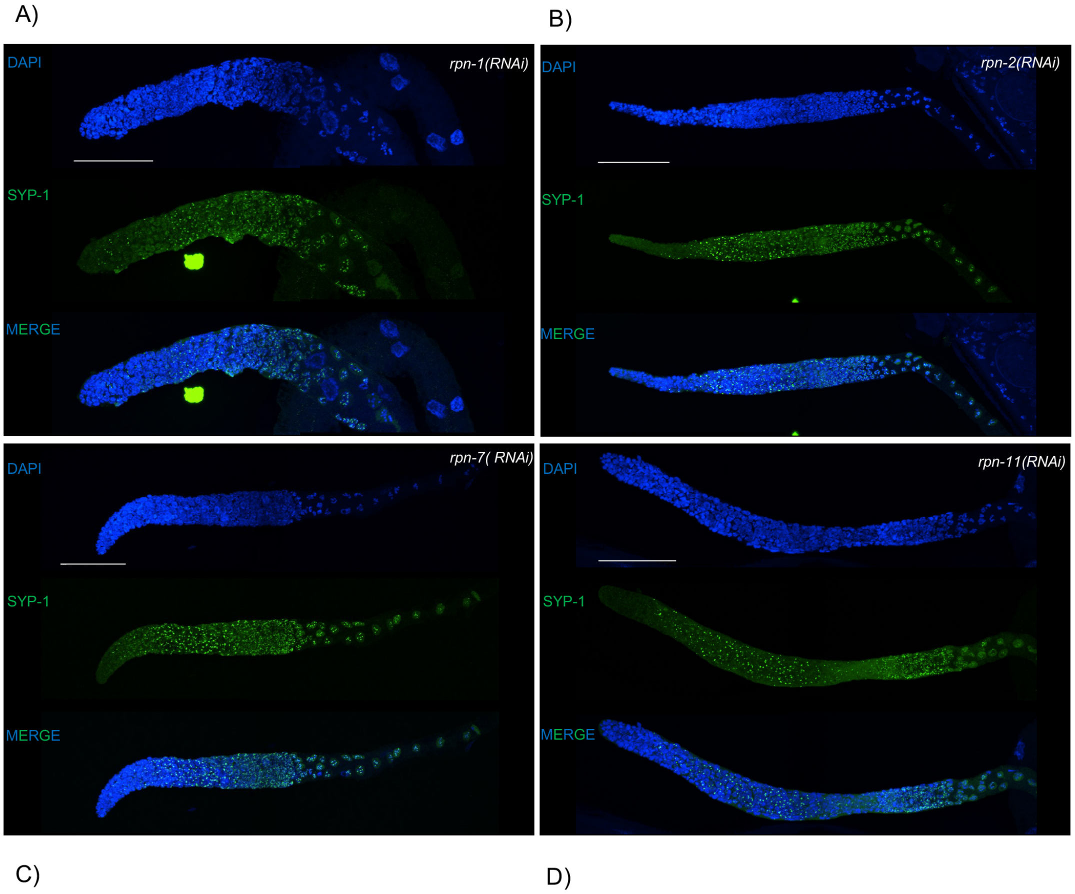


**Supplemental Figure 3: RNAi knockdown of 19S RPN subunits RPN-1, RPN-2, RPN-7, or RPN-11 resulted in a severe SC phenotype.** Extended region of SYP-1 PCs up until mid-pachytene, with almost all nuclei presenting at least one PC and no polymerization of SYP. Premature polarization of SYP-1 was present at late pachytene nuclei. Scale bar = 50 μm.

**Supplemental Figure 4**

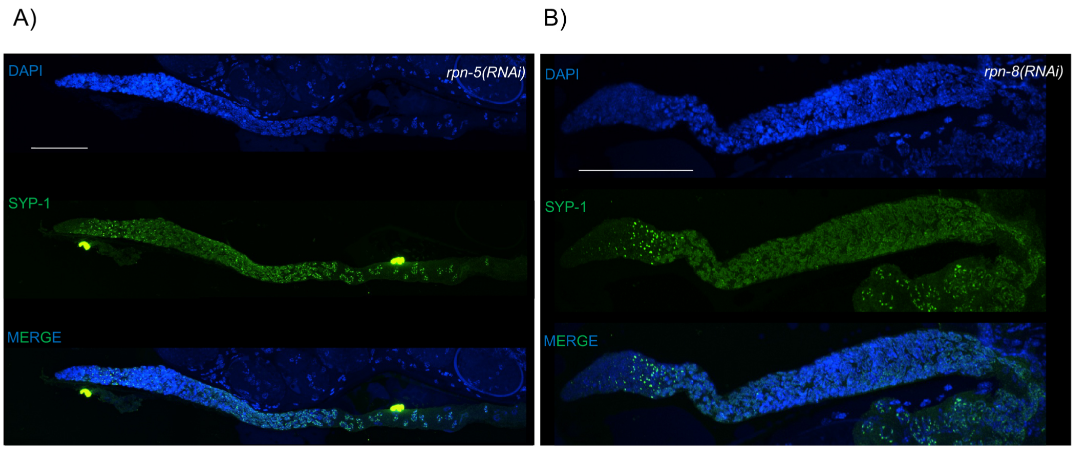


**Supplemental Figure 4: RNAi knockdown of 19S RPN subunits RPN-5 or RPN-8 resulted in a mild SC phenotype.** Extension of SYP-1 PCs region reaching early pachytene, with an abundant number of fully polymerize nuclei in mid-pachytene. Premature polarization is also observed at late pachytene nuclei. Scale bar = 50 μm.

**Supplemental Figure 5**

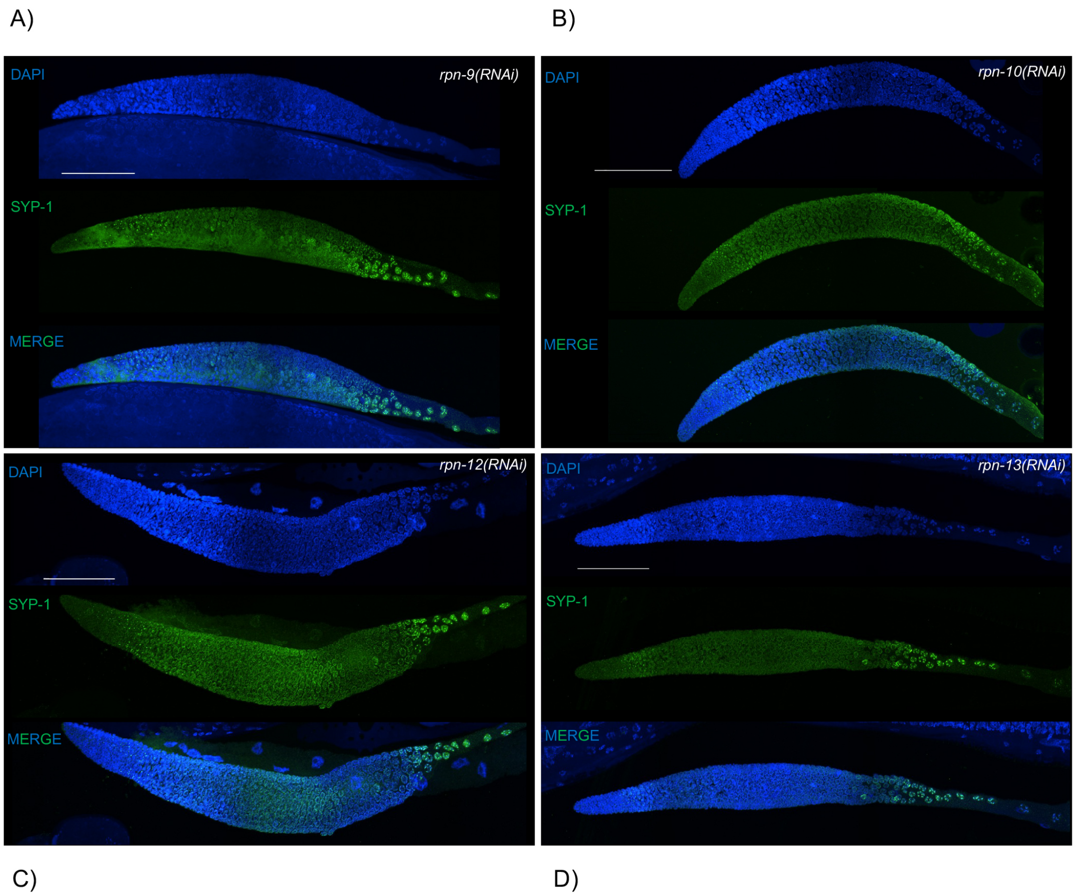


**Supplemental Figure 5: RNAi knockdown of 19S RPN subunits RPN-9, RPN-10, RPN-12 or RPN-13 presented no SC phenotype.** Full polymerization of SYP-1 through pachytene stage and correct timing of polarization to the short arm of the chromosome at diplotene stage comparable to control. Scale bar = 50 μm.

**Supplemental Figure 6**

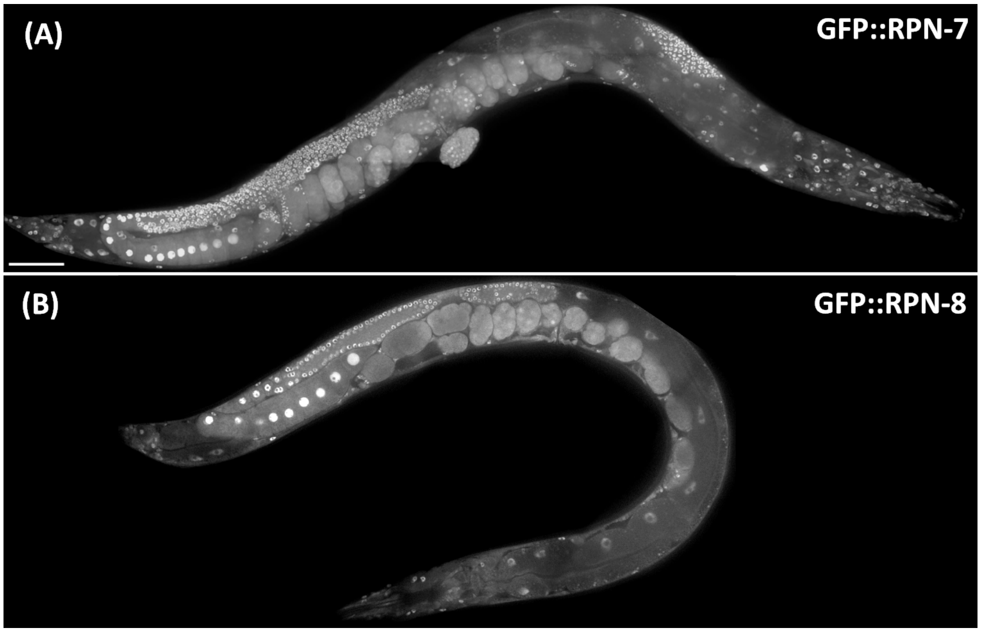


**Supplemental Figure 6. GFP::RPN-7 and GFP::RPN-8 are ubiquitously expressed in the germ line and soma**. Whole worm images of adult hermaphrodites expressing (A) GFP::RPN-7 (Max IP) and (B) GFP::RPN-8 (Single Z stack) exhibiting ubiquitous expression of RPN-7 and RPN-8 in the germ line tissues, soma and embryos with bright nuclear and relatively dim cytoplasmic expression. Scale bar represents 50μm.

**Supplemental Figure 7**

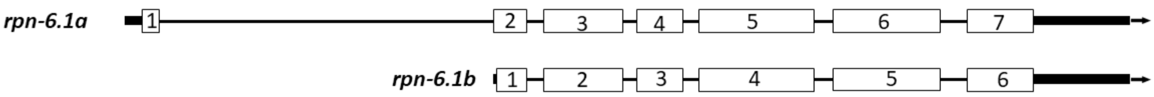


**Supplemental Figure 7: Schematic of the two *C. elegans* protein isoforms for RPN-6.1.** Boxes indicate exons while black lines indicate introns. Thick black lines indicate either 5' or 3' untranslated regions. RPN-6.1 isoform A (RPN-6.1A) contains an extra exon at the C-terminus of the protein, while RPN-6.1 isoform B (RPN-6.1B) is missing that first exon.

**Supplemental Table 1: *C. elegans* proteasome subunits and their human and yeast orthologs.** Includes a list of all the various 19S proteasome subunits in *C. elegans* with their number of predicted splice isoforms, size of the protein in kDA, chromosomal location, and the identified human and yeast ortholog.

| ***C. elegans* proteasome subunit** | **Number of isoforms** | **Size (kDa)** | **Chromo-some** | **Human ortholog** | **Yeast ortholog** |
|---|---|---|---|---|---|
| RPN-1 | 1 | 107kDa | **IV** | PSMD2/S2 | RPN1 |
| RPN-2 | 2 | a- 95.5kDa<br>b- 106kDa | **III** | PSMD1/S1 | RPN2 |
| RPN-3 | 1 | 57.5kDa | **III** | PSMD3/S3 | RPN3 |
| RPN-5 | 1 | 56.4kDa | **II** | PSMD12 | RPN5 |
| RPN-6.1 | 2 | a- 49.1kDa<br>b- 47.1kDa | **III** | PSMD11/S9 | RPN6 |
| RPN-6.2* | 2 | a- 46.8kDa<br>b- 24.2kDa | **III** | PSMD11/S9 | RPN6 |
| RPN-7 | 1 | 47.6kDa | **IV** | PSMD6/S10 | RPN7 |
| RPN-8 | 1 | 40.7kDa | **I** | PSMD7/S12 | RPN8 |
| RPN-9 | 1 | 44.1kDa | **II** | PSMD13/S11 | RPN9 |
| RPN-10 | 1 | 37.3kDa | **I** | PSMD4/S5a | RPN10 |
| RPN-11 | 1 | 34.6kDa | **II** | PSMD14/<br>Poh1/Pad1 | RPN11 |
| RPN-12 | 1 | 28.8kDa | **II** | PSMD8/S14 | RPN12 |
| RPN-13 | 2 | a-39.8kDa<br>b-43.2kDa | **III** | ADRM1 | RPN13 |
| DSS-1 /<br>RPN-15 | 1 | 9.5kDa | **III** | PSMD9/<br>Dss1/ Rpn15 | SEM1 /<br>DSS1 |
| RPT-1 | 1 | 48.6kDa | **V** | PSMC2/S7 | RPT1 |
| RPT-2 | 1 | 49.7kDa | **V** | PSMC1/S1 | RPT2 |
| RPT-3 | 1 | 46.3kDa | **III** | PSMC4/S6 | RPT3 |
| RPT-4 | 2 | a- 45.8kDa<br>b- 44.9kDa | **II** | PSMC6/S10 | RPT4 |
| RPT-5 | 1 | 48.1kDa | **I** | PSMC3/S6a | RPT5 |
| RPT-6 | 1 | 46.2kDa | **III** | PSMC5/S8 | RPT6 |

* RPN-6.2 is a sperm-specific proteasome subunit (personal communication Dr. Lynn Boyd)

**Supplemental Table 2. Strains list**

| Strain name | Description | Genotype | Source |
|---|---|---|---|
| WDC1 | *gfp::rpn-7* | *rpn-7(ana1[gfp::rpn-7])* | This study |
| WDC2 | *gfp::wee-1.3* | *wee-1.3(ana2[gfp::wee-1.3])* | Fernando, *et al.*, 2020 |
| WDC3 | *gfp::rpn-6.1* | *rpn-6.1(ana3[gfp::rpn-6.1])* | This study |
| WDC4 | *gfp::rpn-8* | *rpn-8(ana4[gfp::rpn-8])* | This study |
| WDC5 | *gfp::rpn-9* | *rpn-9(ana5[gfp::rpn-9])* | This study |
| WDC6 | *gfp::rpn-12* | *rpn-12(ana6[gfp::rpn-12])* | Fernando, *et al.,* 2020 |
| WDC12 | *rpn-6.1::OLLAS* | *rpn-6.1(ana12[rpn-6.1::OLLAS])* | This study |
| IT1187 | *UbG76V::GFP::H2B* | *unc-119(ed3) III; kpIs100 [pie-1p::Ub(G76V)::GFP::H2B::drp-1 3' UTR; unc-119(+)]* | Kumar, *et al.,* 2018 |

**Supplemental Table 3. crRNA sequences and properties**

| Gene | Strain | crRNA # | crRNA sequence 5'->3' | %GC | Distance* |
|---|---|---|---|---|---|
| *rpn-6.1* | *gfp::rpn-6.1* | crRNA18 | gtgaactcgtttctctcatt | 40 | 1bp |
| *rpn-7* | *gfp::rpn-7* | crRNA20 | cattttaaggaggatgacag | 40 | 1bp |
| *rpn-8* | *gfp::rpn-8* | crRNA21 | acatctccgtctttgttagt | 40 | 9bp |
| *rpn-9* | *gfp::rpn-9* | crRNA23 | actgctcaagactacctcaa | 45 | 17bp |
| *wee-1.3* | *gfp::wee-1.3* | crRNA29 | gtgaaaatggacgacacaga | 45 | 8bp |
| *rpn-12* | *gfp::rpn-12* | crRNA34 | agccagaagatttttatggg | 48 | 9bp |
| *rpn-6.1* | *rpn-6.1::OLLAS* | crRNA42 | tacaattcatgcaatgggag | 40 | 44bp |

**Supplemental Table 4. Primers to generate repair templates and diagnostics primers**

| Strain | Silent mutations | Primers for repair oligo generation or ssODN and Diagnostics primers (5'→3') |
|---|---|---|
| *gfp::rpn-6.1* | c→g | oAKA334 Fwd:<br>ttcaaaaaattattttaattgacacaacttttcgtgctaatgtccaagggagaggagctctt<br>oAKA335 Rev:<br>atattattagtgtcttctcgtgaactcgtttctctcttgtagagctcgtccattc<br>*Diagnostics*<br>oAKA386 Fwd: gttttgacatcctcgaagctg<br>oAKA387 Rev. cttcgttgtggttaaattgcac |
| *gfp::rpn-7* | g→a | oAKA338 Fwd:<br>atttcaagtgttcatatttcattttaaggaggatgtccaagggagaggagctctt<br>oAKA339 Rev:<br>tcatctacgggtttcttcgtgctctttttggcagcttctgtcttgtagagctcgtccattc<br>*Diagnostics*<br>oAKA388 Fwd: ggccgctttttaacgtttgc<br>oAKA389 Rev. gaacttgcggagtaatccct |
| *gfp::rpn-8* | c→t,<br>a→c,<br>g→t | oAKA340 Fwd:<br>ttgtgataaatttattttcgtttttttagaagaatgtccaagggagaggagctctt<br>oAKA341 Rev:<br>agaacatctacagctttgacagtcgccacatctccatcgttgttggttggagccttgtagagctcgtccattc<br>*Diagnostics*<br>oAKA392 Fwd: gcgtttctcactgttatgtcg<br>oAKA393 Rev: ccatgtcgaggaaccatgta |
| *gfp::rpn-9* | t→c,<br>a→c | oAKA344 Fwd:<br>ctcaatttttaatgtatcgagaatttctcaggatgtccaagggagaggagctctt<br>oAKA345 Rev:<br>ccattggcagccgccagctttccgttgaggtagtcgtgggcggtcttgtagagctcgtccattc<br>*Diagnostics*<br>oAKA394 Fwd: ccatgcgcctttaattgctg<br>oAKA395 Rev: gaaatgcaaatctcgacgagc |
| *gfp::rpn-12* | c→t | oAKA498 Fwd:<br>tatcaaattaaaacattattggatttaagaaaatgtccaagggagaggagctctt<br>oAKA499 Rev:<br>tcctttgcccacacagccagaagatttttatgggcagcagacttgtagagctcgtccattc<br>*Diagnostics*<br>oAKA415 Fwd: accttcacagaatcgtcgag<br>oAKA506 Rev: cattcaaatcgctggaggca |
| *rpn-6.1* | g→a | oAKA456 ssODN |

| | | |
|---|---|---|
| *::ollas* | | gtaccaaactgctctggatacaattcatgcaatgggagaagttgtcgatgcactct atagtaatgcttcgaaaattaactccggattcgccaacgagctcggaccacgtctc atgggaaagtgatgatttcttcgaatcttcatttttgttctgc<br>*Diagnostics*<br>oAKA521 Fwd: aggcgagggaatgcttattg<br>oAKA522 Rev: cgacgtattcctccgtgtat |

20nt crRNA sequences are underlined.
